# MoveFormer: a Transformer-based model for step-selection animal movement modelling

**DOI:** 10.1101/2023.03.05.531080

**Authors:** Ondřej Cífka, Simon Chamaillé-Jammes, Antoine Liutkus

## Abstract

The movement of animals is a central component of their behavioural strategies. Statistical tools for movement data analysis, however, have long been limited, and in particular, unable to account for past movement information except in a very simplified way. In this work, we propose MoveFormer, a new step-based model of movement capable of learning directly from full animal trajectories. While inspired by the classical step-selection framework and previous work on the quantification of uncertainty in movement predictions, MoveFormer also builds upon recent developments in deep learning, such as the Transformer architecture, allowing it to incorporate long temporal contexts. The model predicts an animal’s next movement step given its past movement history, including not only purely positional and temporal information, but also any available environmental covariates such as land cover or temperature. We apply our model to a diverse dataset made up of over 1550 trajectories from over 100 studies, and show how it can be used to gain insights about the importance of the provided context features, including the extent of past movement history. Our software, along with the trained model weights, is released as open source.

## 1 Introduction

The movement of animals is a central component of their behavioural strategies to best exploit the landscape they live in, to find a mate or to avoid predators, for instance. The role that these movements have beyond the individual, for instance in shaping animals’ ecosystem impacts, is clear. Accordingly, and thanks to the technological developments that are allowing to collect more detailed movement data on more individuals and species each day, the study of animal movement has become an important goal of ecology [1].

For a long time, however, statistical tools to analyze movement data were lacking or limited. Over time, though, purely pattern-based descriptions (e.g. home-range analyses) have been complemented by regression models allowing to infer the effects of spatio-temporal features on movement. *Step-selection function* (SSF) models, which compare actual movement steps with realistic candidate ones, are now routinely used to infer and quantify the effect of environmental variables, such as land cover or temperature, on animal trajectories [2–4].

However, an animal’s movement is likely to be driven not only by spatiotemporal environmental features, but also by some internal knowledge and rules that are unobservable directly. The importance of memory and of an animal’s familiarity with places is increasingly recognized [5–7], and familiarity is usually incorporated into SSF models using a *previously visited* yes/no variable, or a *time-spent* variable, often calculated over an arbitrary time window [8, 9]. Memory of places and their characteristics can also lead to routine movement behaviours. Traplining, in which an individual travels to the same places in the same order, is rare, but it is clear from visual inspection of animal trajectories that many animals display some form of routine movement behaviours. But for traplining, which has received a lot of interest, the study of routine movement behaviour has remained extremely limited [10]. Riotte-Lambert et al. [11] showed how conditional entropy, calculated using the information on visits to patches, could be used as a metric of routine in movement. That metric has not been used much since then, possibly because the need to determine sites may render its application difficult on data collected in nature, where patches can be difficult to determine, be diffuse, or not exist at all. Further work is needed to describe and explain routine movements, which result from the interaction between memorized knowledge, movement rules and environmental context. Additionally, we are not aware of any work that has focused on how to incorporate complex information about past movement and environmental context into predictive models of animal movement, although it should, by definition, improve predictions. The question: ‘To what extent past movements inform where an animal is likely to go next?’ remains open.

The classic implementation of the SSF framework appears unsuitable to address this difficult question. We therefore developed a new type of model that we named MoveFormer. We conserved the conceptual attractiveness of SSF, but built on the most recent developments in deep learning to embed the information about current and past animal location, movement and environmental context. Our contribution is threefold. First, we propose a model that learns to best predict the next step of a movement trajectory based on a given context length, i.e. a given time-window of information about the past. Second, the proposed approach is flexible enough to allow each step in the context to be defined not only by the locations of the start and end points, but also by any kind of features that could be relevant, in particular environmental variables. Third, we show how the model can be used to gain insights about the importance of the provided context, both in terms of the extent of the past that it is useful to know, and in terms of what kinds of information are most ecologically relevant to predict an animal’s movement. We demonstrate this by comparing the predictions, via information-theoretic metrics and prediction accuracy, for different context lengths or with randomized features. Model training and analyses are conducted on a dataset made up of over 1550 trajectories from over 100 studies, encompassing various species within mammals, birds and reptiles.

We release the MoveFormer source code,^1^ including code for data pre-processing and evaluation, as well as complete hyperparameter settings and the weights of the trained models. See Section 7 for more information.

## Materials and methods

### 2 Data

In this section, we describe our sources of data, specifically: movement data (trajectories consisting of latitudes, longitudes and timestamps), geospatial variables (associated with locations), and taxonomic classification information (associated with each animal).

### 2.1 GPS location data

Our main source of location data is Movebank^2^ [12], an online repository for animal movement data. The location data in Movebank is presented as latitude/longitude pairs along with UTC timestamps and is grouped into trajectories (*deployments*) and associated with (occasionally missing) metadata such as a taxon name, sex, and date of birth. We used the Movebank API to retrieve data from GPS sensors for all 269 studies that were available^3^ for download under a Creative Commons^4^ license (CC0, CC BY and CC BY-NC), obtaining 13 577 trajectories comprising a total of 197 million observations (location events). We subsampled the trajectories (splitting them into segments when necessary) so that observations occur at midnight and at noon (according to local mean time) with a tolerance of ±3 h and so that the time difference between consecutive observations is 9 to 15 h. We discarded trajectory segments shorter than 120 observations, leaving us with 1440 trajectories from 98 different studies [13–165]. See Table 4 in Supplementary information for the full list of studies and their licenses.

We added data from 4 more studies, collected by one of us (S.C-J). These are GPS data from plains zebras [166] and African elephants [167], collected in Hwange National Park in Zimbabwe, and GPS data from plains zebras and blue wildebeest, collected in Hluhluwe-iMfolozi Park in South Africa and yet unpublished. After subsampling and filtering as in the case of the Movebank data, we obtained 73 trajectories.

See Section 7 for information about data availability.

The final dataset contains about 1 million observations from 1506 individuals, grouped into 1513 trajectories with a median length of 408 observations (corresponding to around 204 days). We performed a train/validation/test split, making sure that 1) the validation and test sections contain only frequent species, and 2) each individual appears in exactly one split.^5^ Table 1 details the amounts of data by section and by taxonomic classification and Fig. 1 shows the geographical distribution.

**Figure 1:**
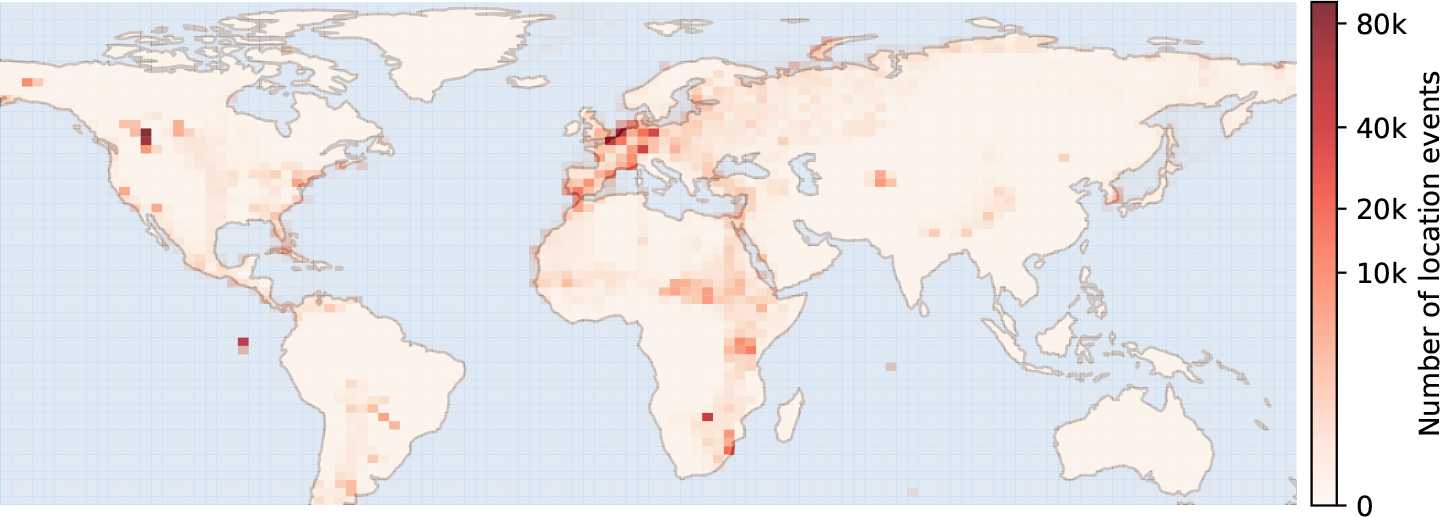
The geographical distribution of the observations in the dataset.

**Table 1:**
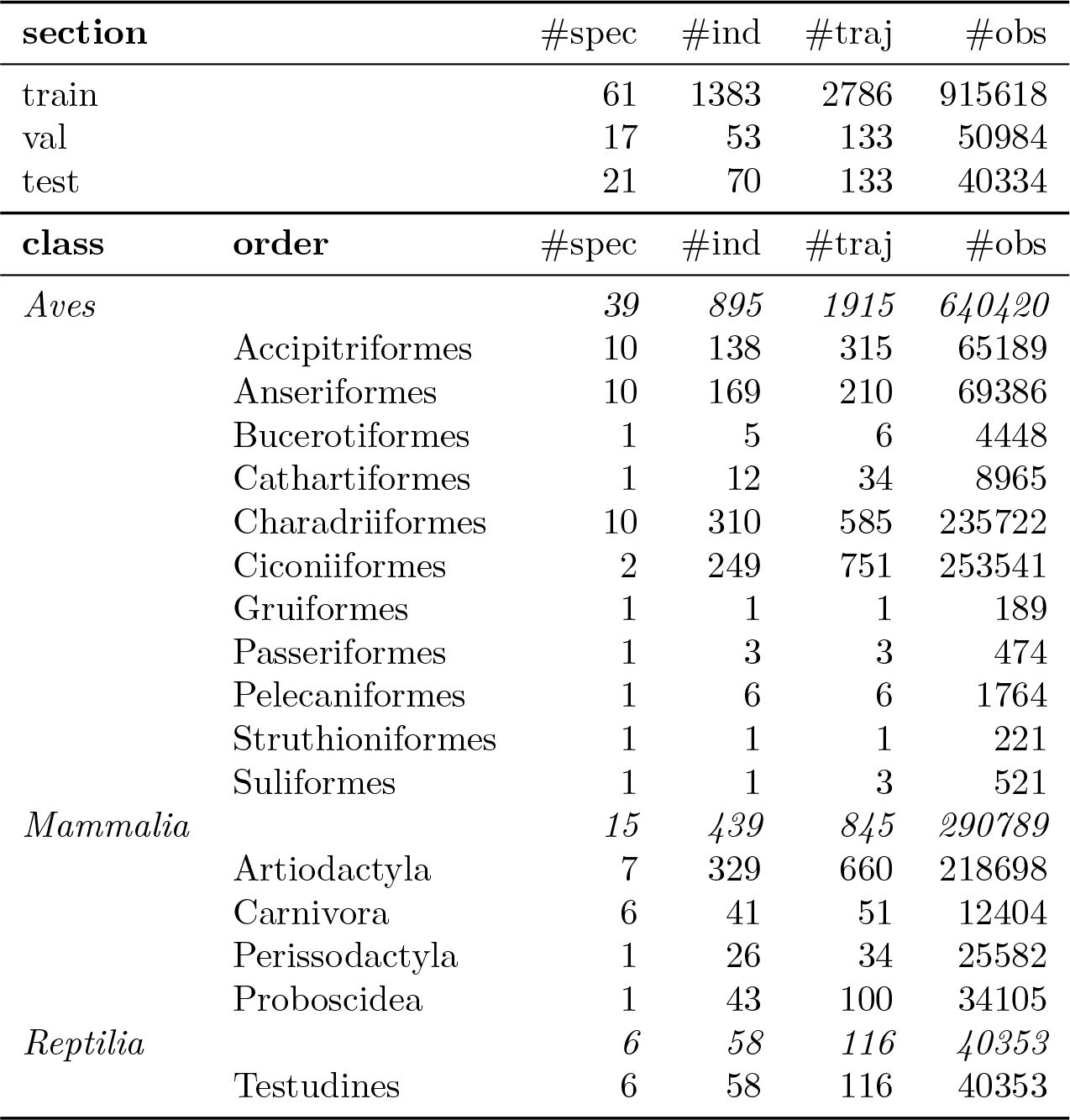
Number of species (#spec), individuals (#ind), trajectories (#traj), and observations (#obs) in the dataset. The upper part of the table displays the counts for each of the sections of the dataset (training, validation, and test). The lower part details the counts for each taxonomic class and order (aggregated over all 3 sections).

| section |  | #spec | #ind | #traj | #obs |
| --- | --- | --- | --- | --- | --- |
| train |  | 61 | 1383 | 2786 | 915618 |
| val |  | 17 | 53 | 133 | 50984 |
| test |  | 21 | 70 | 133 | 40334 |
| class | order | #spec | #ind | #traj | #obs |
| *Aves* |  | 39 | 895 | 1915 | 640420 |
|  | Accipitriformes | 10 | 138 | 315 | 65189 |
|  | Anseriformes | 10 | 169 | 210 | 69386 |
|  | Bucerotiformes | 1 | 5 | 6 | 4448 |
|  | Cathartiformes | 1 | 12 | 34 | 8965 |
|  | Charadriiformes | 10 | 310 | 585 | 235722 |
|  | Ciconiiformes | 2 | 249 | 751 | 253541 |
|  | Gruiformes | 1 | 1 | 1 | 189 |
|  | Passeriformes | 1 | 3 | 3 | 474 |
|  | Pelecaniformes | 1 | 6 | 6 | 1764 |
|  | Struthioniformes | 1 | 1 | 1 | 221 |
|  | Suliformes | 1 | 1 | 3 | 521 |
| *Mammalia* |  | 15 | 439 | 845 | 290789 |
|  | Artiodactyla | 7 | 329 | 660 | 218698 |
|  | Carnivora | 6 | 41 | 51 | 12404 |
|  | Perissodactyla | 1 | 26 | 34 | 25582 |
|  | Proboscidea | 1 | 43 | 100 | 34105 |
| *Reptilia* |  | 6 | 58 | 116 | 40353 |
|  | Testudines | 6 | 58 | 116 | 40353 |

During training and evaluation, we additionally split each trajectory into segments of length *N*_max_ = 500 and subsequently consider each of these segments as a separate trajectory.^6^

### 2.2 Taxon vectors

Each trajectory in our data is associated with a taxon name (most commonly the animal’s species). To obtain a dense vector representation of the taxon, we look up its Wikipedia article and retrieve the associated 100-dimensional embedding vector from Wikipedia2Vec [168].

Wikipedia2Vec embeddings, derived from the text of Wikipedia articles as well as the link structure of Wikipedia, have the property that embeddings of semantically related entities are placed close together in the embedding space. To illustrate that this extends, to some degree, to relationships between biological species, we display in Fig. 2 the PCA (principal component analysis) projections of species embeddings, labeled by higher taxonomic ranks. We also measured the cosine similarity between all pairs of embeddings and found it to be correlated with the number of common ancestors of the two species in the taxonomic hierarchy (Spearman *ρ* = 0.68).

**Figure 2:**
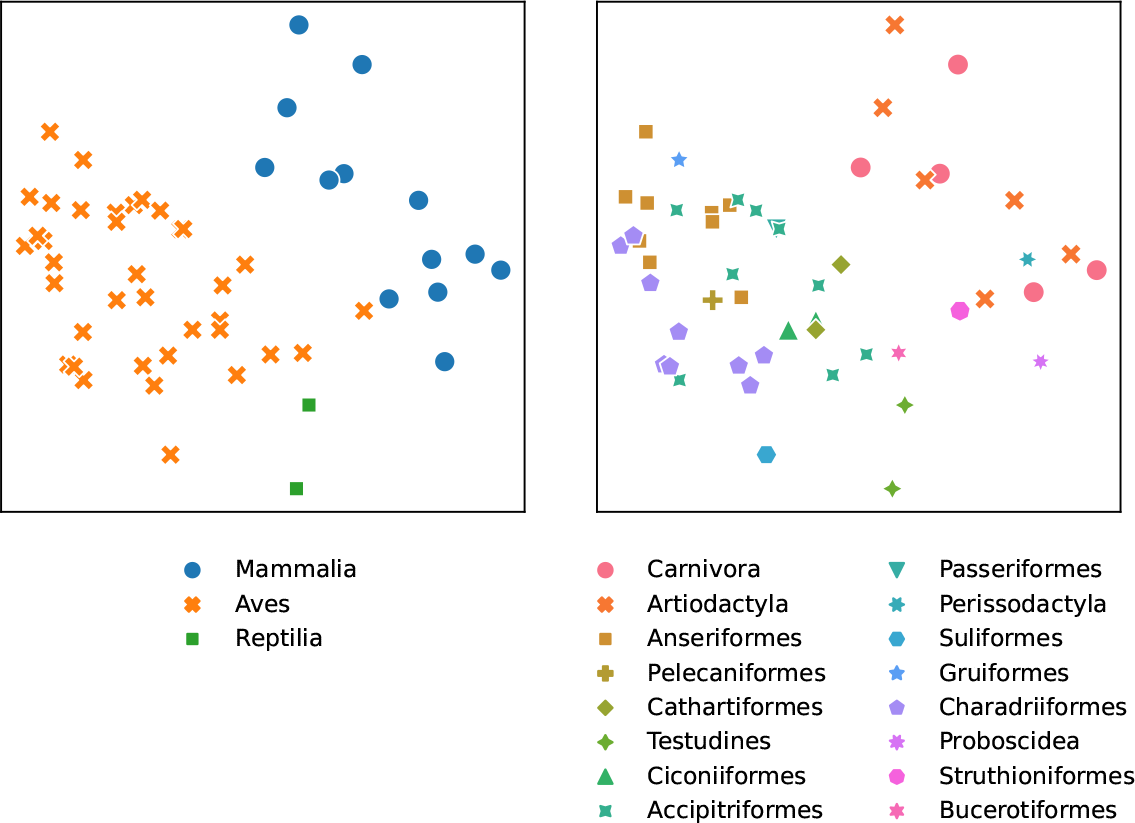
A PCA projection of Wikipedia2Vec embeddings of species, labeled by class (left) and order (right). The *x* and *y* axis correspond to the first two principal components, respectively. Note that since Wikipedia2Vec is a distributed representation, its dimensions are not easily interpretable and have no meaningful units.

Overall, the Wikipedia2Vec embeddings appear to meaningfully encode a species’ position in the phylogeny. Hence, we speculate (though we do not test this in the present work) that their inclusion in the model should help this model to generalize to species that are not present in the training data, at least as long as they are sufficiently similar to those that are.

### 2.3 Geospatial variables

The proposed model is powerful enough to account not only for each trajectory intrinsic dynamics but also for any *third-party* additional information that may be available as covariates. In order to illustrate this, we augment each trajectory data point with exogenous information. For each location, we retrieve the following geospatial variables, which could be ecologically relevant, from publicly available raster data:

- 2009 Human Footprint, 2018 Release [169, 170] (resolution: *∼*1 km); we normalize the values between 0 and 1 and use linear interpolation when retrieving the values by location;
- 19 bioclimatic variables from WorldClim 2.1 [171] (resolution: *∼*1 km), listed in Table 6 in Supplementary information; we standardize the values (zero mean, unit variance) and use nearest neighbor interpolation;
- land cover classification (23 classes) from Copernicus Global Land Service, version 3.0.1, epoch 2015 [172] (resolution: *∼*100 m); we use a one-hot encoding and nearest neighbor interpolation.

## 3 Model

Formally, we consider the dataset as composed of *trajectories*, where a trajectory^7^ *ξ*_1…*N*_ of length *N* consists of locations *x*_1…*N*_, corresponding to timestamps *t*_1…*N*_, and any associated variables *z*_1…*N*_, i.e. *ξ*_*n*_ = (*x*_*n*_, *z*_*n*_, *t*_*n*_) as described above. Our main goal is to estimate a model for the next-step prediction task, i.e. for any given *n ∈ {*1, …, *N}*, predict the next location *x*_*n*+1_ from the trajectory prefix *ξ*_1…*n*_ and the next timestamp *t*_*n*+1_.

As a fundamental use case, we are interested in analyzing the effect of available past context on the prediction of *x*_*n*+1_. Specifically, for a varying *context length c∈ {*1, …, *c*_max_*}* (where *c*_max_ is an arbitrary constant), we wish to study the behavior of the prediction of *x*_*n*+1_ given *ξ*_*n*−*c*+1…*n*_ and *t*_*n*+1_. Hence, we are in fact interested in a model accepting as input any *trajectory segment* of length at most *c*_max_, and predicting the next location.

We adopt a step-selection function modelling approach [2, 4], based on selecting the end-point location of a step from a set of candidates. Specifically, for a position *n*+1 within a trajectory, given an associated timestamp *t*_*n*+1_, a set of candidate locations 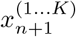 and associated variables 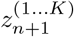, we are interested in estimating a probability distribution over the candidates:

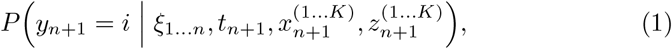

where *i ∈ {*1, …, *K}*.

We propose to model this distribution using a deep neural network, consisting of a *Transformer* [173] *encoder* and a *candidate selection module*, as depicted in Fig. 3. The role of the Transformer is to encode the trajectory up until position *n*, i.e. *ξ*_1…*n*_ along with the timestamp for the next observation, *t*_*n*+1_. The candidate selection module then encodes each candidate 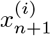 and employs an attention mechanism to compute a probability distribution over the candidates. The model is described in detail in Section 3.1, followed by our choice of input representation in Section 3.2.

**Figure 3.**
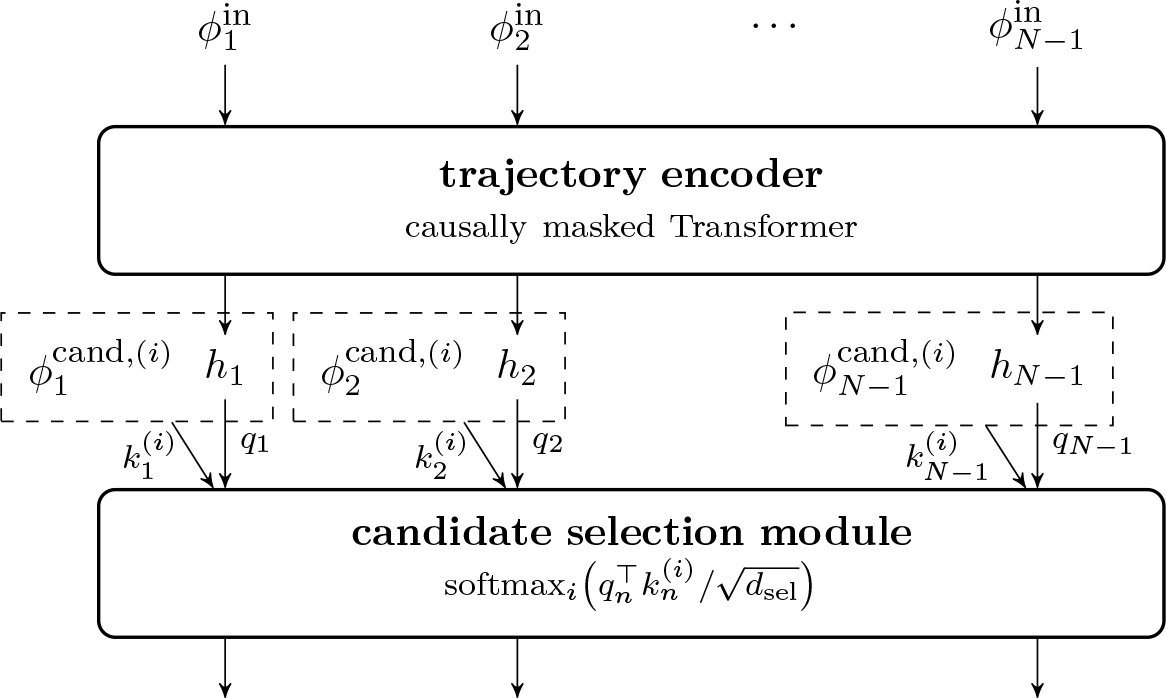
The high-level architecture of MoveFormer. The input to the trajectory encoder is a sequence of embedding vectors 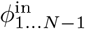, each corresponding to a different data point (location-timestamp pair) in the trajectory. The encoder outputs a sequence of vectors *h*_1…*N−*1_; the causal masking in the encoder causes each *h*_*n*_ to encode only the inputs up to position *n*, i.e. 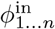. This representation is then fed to the candidate selection module, which uses it as *queries* in an attention mechanism that assigns probabilities to different candidate locations.Both the input embeddings 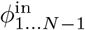 and the candidate embeddings 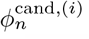 are computed through embedding layers which are not displayed here but described in Section 3.1.1.

**Figure 4:**
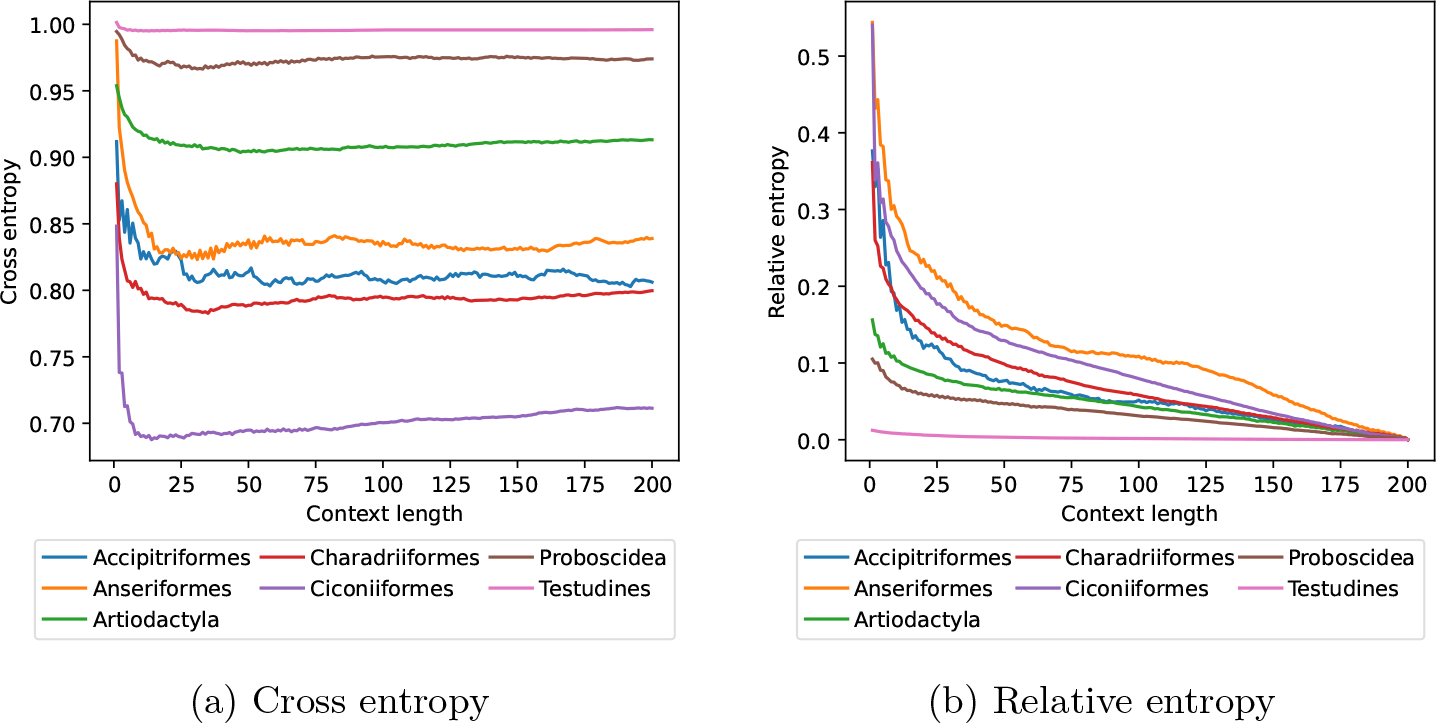
Metric value averages by context length and taxonomic order, computed on the test set (only positions *n > c*_max_ = 200).

In order to train and evaluate this model, we also need a way to generate suitable candidate locations 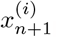. We use a simple but general method employing quantile-based modelling of turning angles and movement distances, as detailed in Section 3.3.

### 3.1 Step-selection model

#### 3.1.1 Input embeddings

We build two sets of embeddings 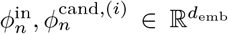, *n ∈ {*1, …, *N −* 1*},i ∈ {*1, …, *K}* such that:

- 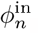, input for the trajectory encoder, depends on *x*_*n−*1_, *x*_*n*_, *t*_*n*_, *t*_*n*+1_, *z*_*n*_;
- 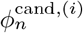, input for the candidate selection module, depends on *x*_*n*_, 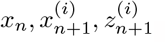.

The inputs are represented as collections of carefully engineered continuous and discrete features that we will describe later (see Section 3.2). Missing (NaN) values are replaced with a special embedding vector learned as an additional parameter. In each case, we project each feature vector to a common embedding space 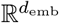, then linearly combine them (with different learnable coefficients in each of the two cases).

More precisely, for *ϕ*^in^:

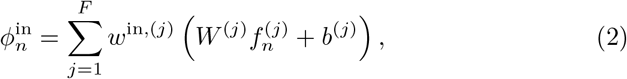

where 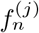 is the *j*-th out of all *F* feature vectors at step *n*, and the learnable parameters are coefficients *w*^in,(*j*)^ ∈ ℝ(we set *w*^in,(*j*)^ = 0 for features we do not wish to consider), biases 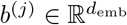 and weight matrices 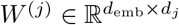. The formula for 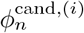 is analogous. As can be seen from Eq. (2), the chosen method for constructing input embeddings allows features to have different dimensions and automatically projects them to the desired embedding dimension (via *W* ^(*j*)^ and *b*^(*j*)^) before applying scaling through *w*^in,(*j*)^.

#### 3.1.2 Trajectory encoder

The trajectory encoder is a Transformer encoder with causally masked attention. It receives the embedding sequence 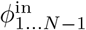and outputs a sequence of vectors *h*_1…*N* −1_ where *h*_*n*_ is a representation of *ξ*_1…*n*_ . The encoder does not use any positional encoding in the conventional sense (encoding the indices 1, …, *N* − 1, as is commonly done in Transformers), but position information is conveyed by the feature representations of the timestamps *t*_1…*N*−1_.

#### 3.1.3 Candidate selection

The candidate selection module is used to select the next location out of a list of candidates. We build upon the common approach that models the probability of an individual being present at a given candidate location via conditional logistic regression [3]; expressed in our notation:

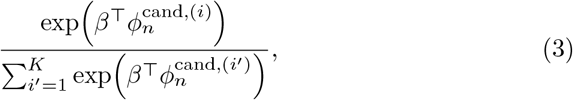

where *β* is a parameter vector.

In this work, in order to incorporate the context representation computed by the trajectory encoder, we replace the global parameter vector *β* with a context-dependent *query vector* 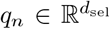, which is a linear projection of the trajectory encoder output *h*_*n*_. We also do not use the raw candidate features 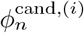 but replace them with a *key vector* 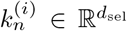, which is computed by concatenating the feature vector with the corresponding encoder output *h*_*n*_ and passing the result through a *candidate encoder* (a fully-connected network): 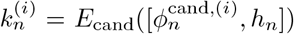. Thus, we arrive at a *dot-product attention mechanism*; scaling the dot products by 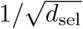 as in Transformer attention [173], we have:

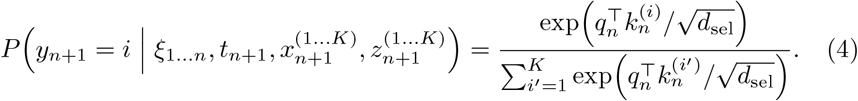

During training, the first candidate location ^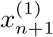^ is taken as the true next location *x*_*n*+1_; the rest of the candidates are randomly sampled around the current location *x*_*n*_ (we detail this process below). This allows us to define a cross entropy loss, which we minimize through stochastic gradient descent using the Adam optimizer:

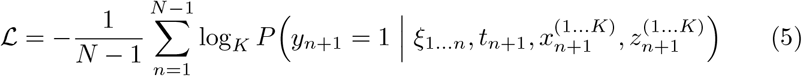

#### 3.1.4 Variable receptive field training

As mentioned above, we aim to evaluate our model on arbitrary trajectory segments up to some maximum length *c*_max_ (this procedure is detailed below in Section 4.1). As can be seen from Eq. (5), our model is effectively being simultaneously trained on all prefixes of the trajectory *ξ*_1…*N*_ . Hence, the model is able to accept segments of variable length as desired, but being only trained on trajectory prefixes may bias it, leading to incorrect predictions on segments that are not prefixes. To alleviate this, we propose a training scheme that intervenes on the attention weights to randomly vary the past context available for each prediction.

In each training batch, we sample a random integer *B* uniformly from {1, …, *N*_max_} and apply a block-diagonal attention mask to the attention matrix (on top of the causal mask) with blocks of size *B* (with the last block truncated if *B* ∤ *N*). As a result, the ranges of positions {1, …, *B*}, {*B* + 1, …, 2*B*}, etc. are prevented from attending to each other, and the corresponding segments are therefore effectively considered as separate trajectories.

### 3.2 Data representation

Let us now describe the feature mappings used for location and time, as well as associated features.

#### 3.2.1 Location

In the raw data, each location *x*_*n*_ is represented as a GPS coordinate pair (latitude, longitude). We represent it as a geodetic normal vector (*n-vector*) *v*(*x*_*n*_) ∈ ℝ^3^.

Additionally, we encode the position relative to the previous location *x*_*n*_ − _1_ a *movement vector μ*(*x*_*n*_ − _1_, *x*_*n*_) ∈ ℝ^2^, obtained by computing the bearing and distance from *x*_*n*_ − _1_ to *x*_*n*_ and converting them to cartesian coordinates. We apply scaling to make the overall root-mean-square (RMS) of the norms of movement vectors computed on the training dataset equal to 1.

Analogously, we encode each candidate location 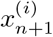 as an n-vector 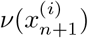 and as a movement vector 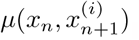

#### 3.2.2 Time

We encode a timestamp *t*_*n*_ as:

- a 10-dimensional vector of sines and cosines with a period of 1 second, 1 minute, 1 hour, 1 day, and 1 tropical (solar) year, respectively, such that their phase synchronizes on January 1st, 2000, at 00:00:00 UTC;
- sin(LMT*/*24 ·2*π*) and cos(LMT*/*24 ·2*π*), where LMT is the local mean time (i.e. UTC adjusted by longitude) in (fractional) hours;
- integer values (one-hot-encoded) representing the calendar month (0– 11), the day of the month (0–30), and the day of the week (0–6) in UTC.

We also encode the time difference w.r.t. the next timestamp *t*_*n*+1_ as a 12-dimensional vector of sines and cosines with the same periods as above, plus a period of 25 years.

While this multi-scale encoding may not be necessary in our case (where the time differences are between 9 and 15 h), we propose it as a generic representation suitable for any time scale from seconds to years (and hence for virtually all existing animal movement data).

#### 3.2.3 Associated variables

For each input and candidate location, we retrieve and pre-process geospatial variables as described in Section 2.3. We also include the taxon vectors (as described in Section 2.2) as an additional encoder feature vector for every element of the input sequence.

### 3.3 Candidate sampling

We sample each candidate location 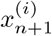 as follows:

- we estimate the current bearing *β* of the animal from the positions *x*_*n*_ and *xn*−1;
- we independently sample a *turning angle* 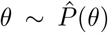 and a *log-distance* 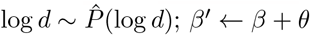;
- we compute 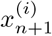 by moving *x*_*n*_ according to *β*^*′*^ and *d*.

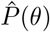 and 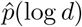 are estimated on the training set as follows:

- We collect all turning angles from the training set and compute the quantiles (estimated using linear interpolation) at 101 equally spaced points 0 = *q*_0_, *q*_1_, …, *q*_100_ = 1. We use them to construct the quantile function of 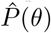 as a piecewise linear function with knots at *q*_0_, *q*_1_, …, *q*_100_.
- We collect the natural logarithms of all non-zero distances between consecutive points in the dataset; we construct the quantile function of 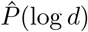 analogously.

We sample from each distribution by drawing a sample from 𝒰 [0, 1] and passing it through the estimated quantile function; this is sometimes called the *increasing rearrangement* [174].

In our experiments, we condition the distributions on the taxon, i.e. we estimate a separate pair of distributions on the section of the training dataset corresponding to each taxon.

### 3.4 Implementation details and hyperparameters

Our implementation of MoveFormer, available as open source software, is written in Python using the PyTorch framework^8^ and the `x-transformers`^9^ package. The code for efficient geospatial variable loading relies on the rasterio^10^ library and is released as a separate package, `gps2var`. See Section 7 for information about code availability.

The trajectory encoder is a 6-layer Transformer with 8 attention heads per layer and a feature dimension of 128. The candidate encoder is a fully-connected neural network with one hidden layer of size 256 and a GELU activation [175]. The candidate selection module has *d*_sel_ = 128. The total number of parameters of the model is around 2.6 million – several orders of magnitude smaller than current state-of-the-art Transformer language models, for instance, but appropriate for the limited-size dataset that we are working with.

The Adam optimizer uses a learning rate of 5 × 10−5 with linear warm-up and exponential decay. We train for 180 epochs with a batch size of 24, taking 7.5 h on a Tesla V100 GPU (note that GPU utilization was only about 20 % and the performance bottleneck appeared to be the geospatial variable loading). We validate on the validation set twice per epoch and use the checkpoint with the lowest validation loss.

The complete hyperparameter settings are included with the source code.

## 4 Analysis methods

### 4.1 Context length analysis

Riotte-Lambert et al. [11] propose to use *conditional entropy* as a measure of uncertainty in predicting the next location given the *c* previous locations. Specifically, given a distribution *P* over sequences of locations, conditional entropy of order *c* can be written as

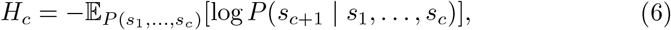

where *P* (*s*_1_, …, *s*_*c*_) is understood as the probability of *c* consecutive locations in a sequence being equal to *s*_1_, …, *s*_*c*_, and *P* (*s*_*c*+1_ | *s*_1_, …, *s*_*c*_) as the conditional probability of *s*_*c*+1_ immediately following the sequence *s*_1_, …, *s*_*c*_. Considering this uncertainty measure as a function of the context length *c*, it may be used to study routine movement behavior.

Riotte-Lambert et al. [11] work with a finite set of discrete locations, allowing them to evaluate the expression (6) empirically on a given trajectory. However, the probability estimates quickly become unreliable with increasing *c* due to data sparsity. Moreover, the method is inapplicable when locations are unique, as in our case.

We propose an alternative way, which is to approximate log *P* using a suitable machine learning model (e.g. our proposed step selection model), so that Eq. (6) becomes *cross entropy* computed on trajectory *segments* of appropriate length. In our case:^11^

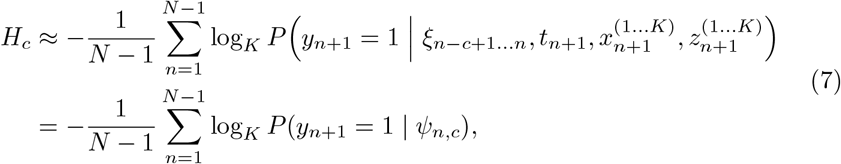

where we collapse all the conditioning variables to *ψ*_*n,c*_ for brevity. For more fine-grained analysis, we may be interested not in the sequence-level cross entropy, but rather in the ‘pointwise’ values, i.e. −log_*K*_ *P* (*y*_*n*+1_ = 1 |*ψ*_*n,c*_).

More generally, we may alternatively choose to examine any metric that can be computed from the probabilities. We adopt the *relative entropy* (also known as the *Kullback-Leibler divergence*) of the prediction with the maximum context length *c*_max_ with respect to the one at context length *c* (as proposed by Cífka and Liutkus [176] in the context of causal language models for text):

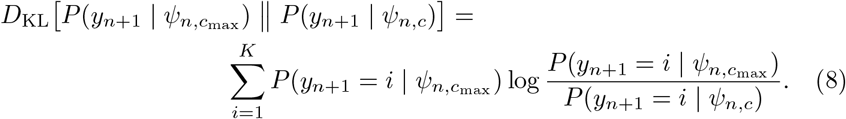

Note that this metric does not depend on the ground truth location, but measures the amount of information gained by considering the maximal context instead of the limited one.

#### 4.1.1 Relevant context length

We may expect that there would be a critical context length *C* after which the above metrics stop improving, as further extending the context does not result in significant information gain. Similarly to Riotte-Lambert et al. [11], we define the *relevant context length C*_*m*_ – for a given metric *m* – as the smallest context length for which the metric reaches its optimum, with a 5% tolerance for robustness to noise:

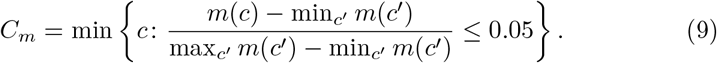

#### 4.1.2 Efficient evaluation

We now discuss how to efficiently compute the probabilities needed to calculate the above metrics, following the procedure proposed for causal language models by Cífka and Liutkus [176]. We may collect all the probabilities in a tensor 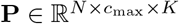 such that

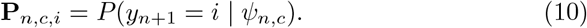

Observe that by running the model on a segment of the trajectory corresponding to indices *n*, …, *n*+*c*_max_ −1 for a given *n*, we obtain all the values **P**_*n*+*c*_ −_1,*c*, ∗_ for *c* ∈{1, …, *c*_max_} . We may also notice that 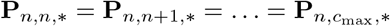 for any *n < c*_max_. Hence, we can efficiently fill in the tensor **P** using *N* runs of the model on segments of length at most *c*_max_.

### 4.2 Candidate feature importance

While the parameters of step-selection models fitted by conditional logistic regressions or point-process models are directly interpretable [4], deep learning models are known as ‘black boxes’ that require special techniques to be interpreted post-hoc. A simple but popular technique [177, 178] is based on testing the model on a dataset with the values of a given feature randomly permuted. While aware of the caveats related to using this technique with correlated features [179], we employ it here to demonstrate the possibility of interpretation, and leave more advanced techniques for future work.

Specifically, we study how individual *candidate features* (components of 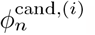) influence the selection of candidates. We pick a feature (or a group of features), and for every observation in the dataset, we randomly shuffle the feature’s values among the *K* candidates (in contrast to Fisher et al. [178], who shuffle values across the entire dataset). The aim is to make the feature completely uninformative while maintaining its values plausible in the given context. We evaluate the model on both the permuted and the original dataset, and use the difference in performance as a measure of the importance of the selected feature.

## Results and discussion

### 5 Results

#### 5.1 Validation

We evaluate the proposed model (here dubbed VarCtx) against variants to serve as baselines:

- FullCtx is a variant without the variable receptive field training (see Section 3.1.4);
- NoAtt is a model where all the attention layers are removed from the Transformer encoder, so that information is not allowed to flow between different positions in the sequence;
- NoEnc is a model where the Tranformer encoder is removed, i.e. we have 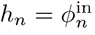

Note that the last two variants have a receptive field of 1 (i.e. only the features at position *n* are available for predicting the location at *n* + 1). To simulate this for VarCtx and FullCtx in a comparable way, we test these in a regime (denoted by +diag) where the attention matrices are restricted to an identity matrix, i.e. each position can only attend to itself.

After running each of the above models on the test set, we compute the following metrics:

- xent@16: cross entropy (Eq. (5)) computed with 16 candidates;
- xent@100: cross entropy computed with 100 candidates;
- acc 1/16: accuracy (i.e. how often the top scoring candidate is the ground truth) with 16 candidates,
- acc 10/100: top-10 accuracy (i.e. how often the ground truth is among the 10 top scoring candidates) with 100 candidates.

The results, averaged over all trajectories, are presented in Table 2. We note that the results are very consistent across all metrics, and we found all pairs of metrics to be strongly correlated (Pearson *ρ >* 0.87, computed over all models and trajectories).

**Table 2:** Results for different variants of the model. FullCtx: trained on full trajectories (max. length 500); VarCtx: trained with variable receptive field; NoAtt: no attention layers; NoEnc: no encoder; diag: attention restricted to diagonal matrix during inference. Xent: cross entropy (lower is better), acc: accuracy (higher is better).

| model | xent@16 ↓ | xent@100 ↓ | acc 1/16 ↑ | acc 10/100 ↑ |
| --- | --- | --- | --- | --- |
| FULLCTX | 0.869 | 0.909 | 0.198 | 0.293 |
| FULLCTX+DIAG | 0.990 | 0.998 | 0.102 | 0.157 |
| VARCTX | **0.847** | **0.894** | **0.221** | **0.323** |
| VARCTX+DIAG | 0.932 | 0.954 | 0.136 | 0.204 |
| NOATT | 0.919 | 0.945 | 0.157 | 0.231 |
| NOENC | 0.928 | 0.950 | 0.148 | 0.217 |

Both FullCtx and VarCtx outperform the rest of the models, which have a receptive field length of 1. This is evidence that providing past movement as context is beneficial. Interestingly, VarCtx yields better results than FullCtx, possibly because the variable receptive field training scheme effectively makes the training data more diverse, alleviating overfitting.

We can also observe that the results of VarCtx+diag are closest to those of the models trained with minimum context (NoAtt, NoEnc). This suggests that the performance of VarCtx is not strongly degraded by limiting its receptive field at test time (unlike that of FullCtx), validating our variable receptive field training approach.

Finally, we noticed large performance differences between species. For the VarCtx model, we calculated the average cross entropy for each taxonomic order (see Table 3) and found that it tends to be lower (i.e. better) for orders with a higher number of observations in the training set (Pearson *ρ* = −0.71).

**Table 3:** VarCtx validation cross entropies by taxonomic order, along with numbers of observations in the training data.

| class | order | xent@16 | #train |
| --- | --- | --- | --- |
| Aves | Accipitriformes | 0.815 | 58 994 |
|  | Anseriformes | 0.827 | 64 008 |
|  | Cathartiformes | 1.057 | 8653 |
|  | Charadriiformes | 0.815 | 205 602 |
|  | Ciconiiformes | 0.697 | 237 304 |
| Mammalia | Artiodactyla | 0.928 | 201 464 |
|  | Carnivora | 0.986 | 12 282 |
|  | Proboscidea | 0.980 | 24 870 |
| Reptilia | Testudines | 0.998 | 34 577 |

### 5.2 Context length analysis

In this section, we demonstrate the ability to use the VarCtx to study the dependence of the predictions on the length *c* of the available past context, as described in Section 4.1. We set *c*_max_ = 200 and *K* = 16.

First, we display in Fig. 5 the average cross entropy and relative entropy as a function of context length and by taxonomic order, and in Fig. 5 examples for concrete observations, with the relevant context length *C* highlighted. We observe that the best predictions tend to be achieved around context lengths 10–50, which corresponds to 5–25 days.

**Figure 5:**
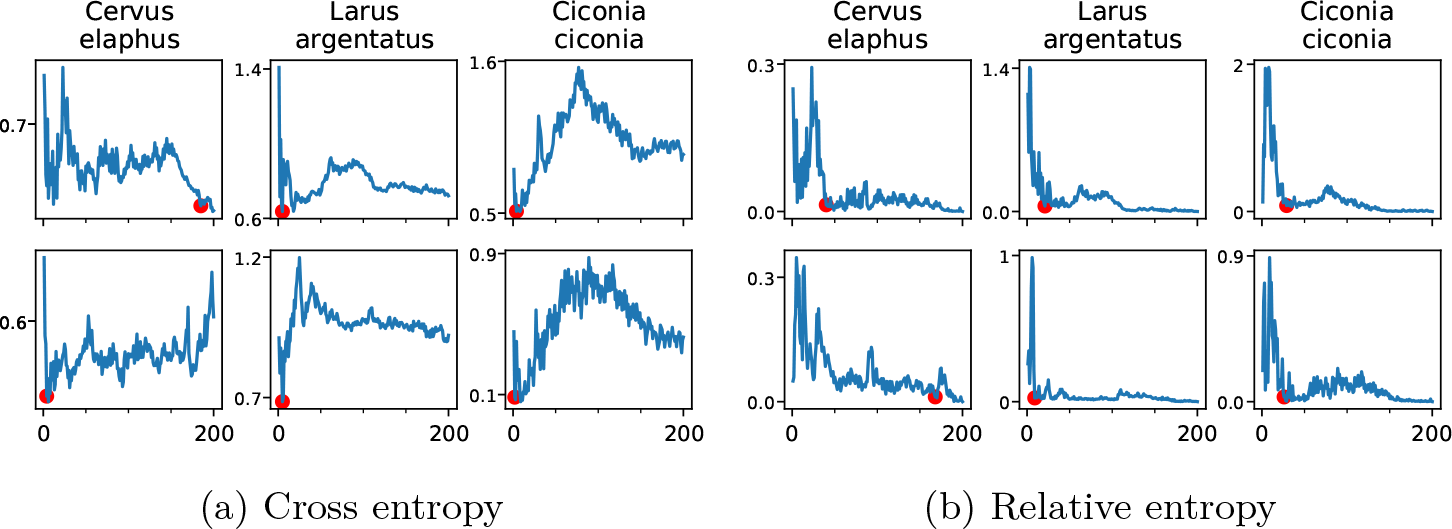
Examples of metric values (pointwise, i.e. for a single observation within a given trajectory) plotted as a function of context length. Top and bottom correspond to different (random) positions within the same trajectory. The red dot marks the relevant context length (where the metric reaches 5 % of its min-max range).

Apart from the clear inter-species differences in cross entropy already noted in the previous section (Table 3), we also observe some differences in *relative* entropy, though less marked. For example, while *Ciconiiformes’* movements are substantially easier (in terms of cross entropy) for our model to predict than those of *Anseriformes*, both have a similar relative entropy profile, indicating that the amount of information contributed by each time scale is similar for both taxa. On the other hand, note that the flat relative entropy profile of *Testudines* simply reflects a failure of our model to accurately predict their movements at any time scale – as evidenced by the cross entropy values being close to 1 –, which is possibly due to an insufficient amount of reptile training data.

Fig. 6 shows the distribution of the relevant context length *C* for each taxon in the test set. There are apparent differences between taxa, but we also note the large variability *within* each taxon that could be of interest in itself.

**Figure 6.**
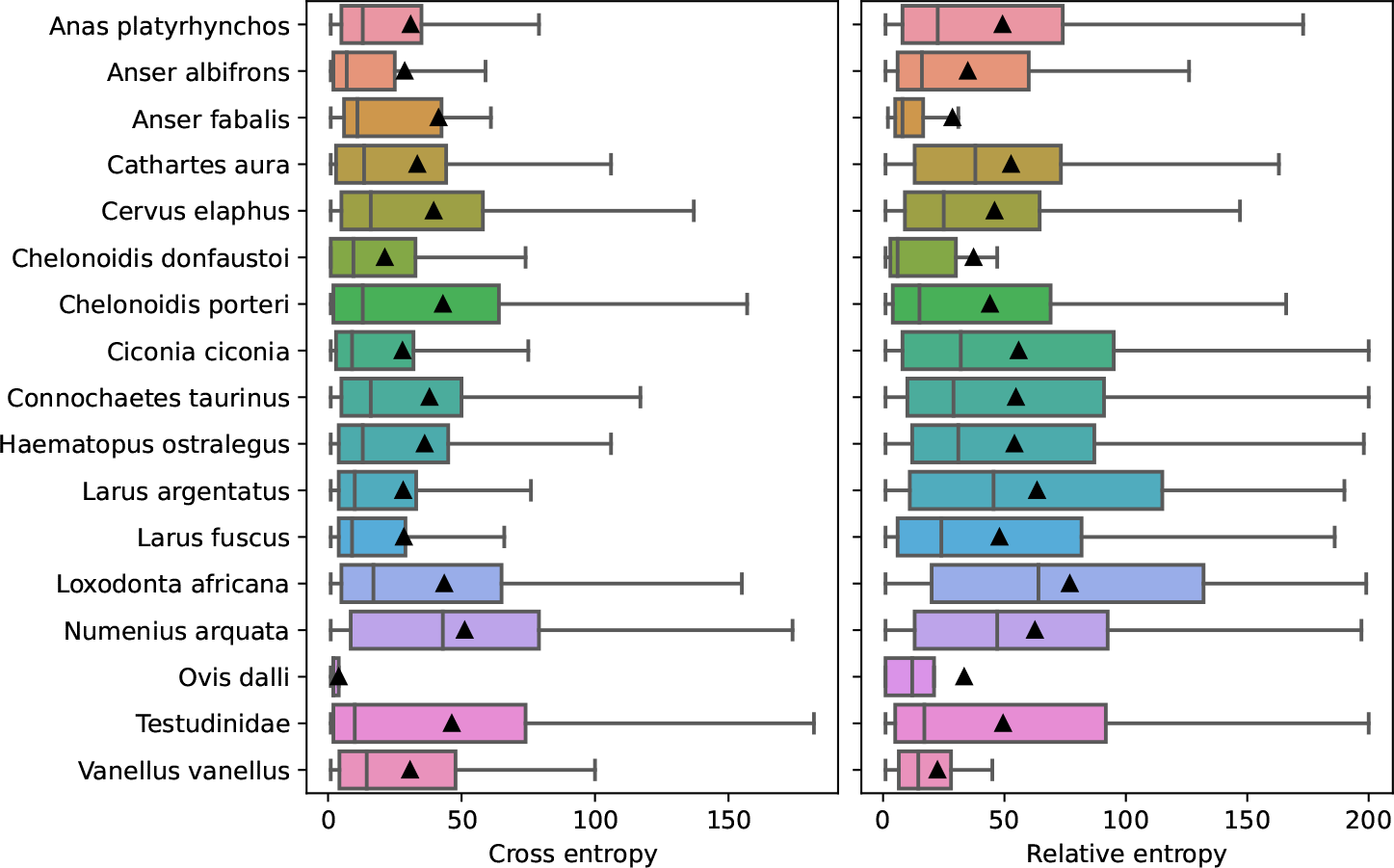
Relevant context length *C* by taxon, computed using cross entropy and relative entropy (pointwise values, as shown in Fig. 5), respectively. The black triangles indicate means.

### 5.3 Candidate feature importance

We present in Fig. 7 the results of the feature importance experiment. Vector features (location) are treated as groups; bioclimatic variables are tested both individually and as a group.

**Figure 7.**
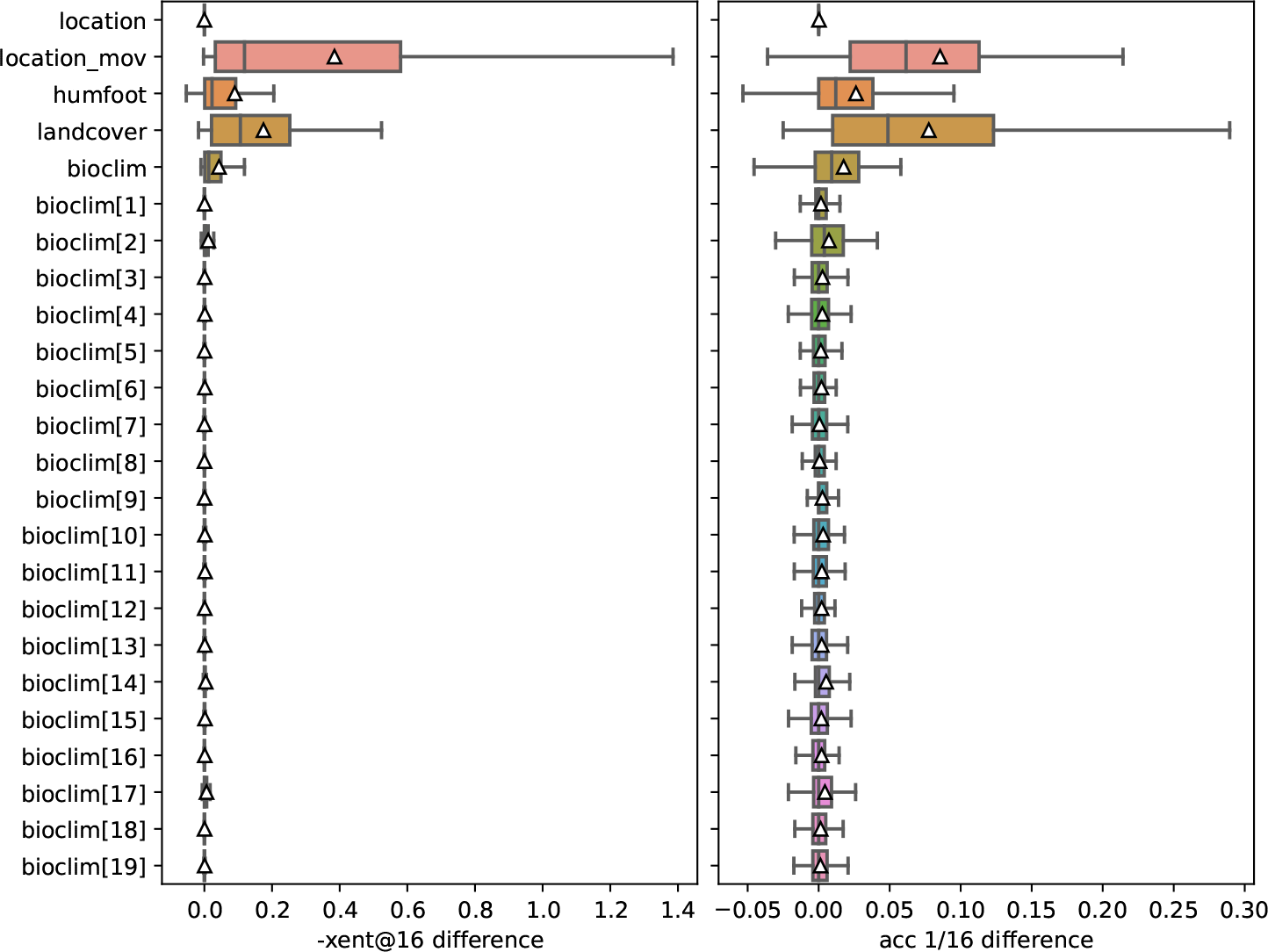
Candidate feature importances computed as differences between performance (measured using negative cross entropy and accuracy with 16 candidates) with and without permuted feature values. The plot shows the distribution over test trajectories. The features are, from top to bottom: location n-vector, movement vector, human footprint, land cover, and finally the 19 bio-climatic variables (BIO1 to BIO19; see Table 6 in Supplementary information), first as a group and then each individually.

The most important features found by this method are movement vector and land cover, followed by human footprint. The bioclimatic variables appear to have relatively low impact, with the most important ones being BIO2 (mean diurnal range), BIO14 and BIO17 (both related to precipitation). Interestingly, global location (represented as n-vectors) seems to be the least important feature, possibly because it is difficult to exploit for candidate selection compared to the relative location information provided by the movement vectors.

Note that only *candidate* features 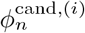 are tested here, and the results do not say anything about the *input* (past observation) features 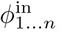 . For example, global location, which we found to be unimportant as a candidate feature, may well turn out to be an important past context feature.

## 6. Discussion

In this work, we propose a new model to learn from animal trajectories. Inspired by the classical step-selection framework [2] and previous work on the quantification of uncertainty in movement predictions [11], we designed MoveFormer, a step-based model of movement that builds upon recent developments in deep learning, such as the Transformer architecture. This allowed us to meet our initial goal to endow the model with a unique ability to learn how past movements influence current and future ones. Although this is an important question in movement ecology, it has remained poorly addressed so far because classical step-selection functions or other movements models are unable to account for past information except in a very simplified way (e.g. by including a feature indicating whether or not the animal has previously visited a given site).

An important contribution of this work is also to generalize the suggestion of Riotte-Lambert et al. [11] to use conditional entropy calculated over visits to discrete sites as a way to measure movement uncertainty. Although attractive, the difficulty of discretizing trajectories to meaningful ‘sites’ has slowed down the application of this idea. Here, we extend it to locations acquired in continuous space and propose cross-entropy and relative entropy, estimated through the movement model, as a more general approach. This allows to estimate the *relevant context length* (‘relevant order of dependency’ in Riotte-Lambert et al. [11]), i.e. the amount of the past that significantly improves the predictions about further movements. We did so in this study, and to the best of our knowledge, our study therefore provides the first estimation of how much of the past one needs to know to improve predictions of animal movements.

Our results suggest that for most datasets, predictions are improved when integrating the information from about a few days to two or three weeks before the movement to be predicted. Why this is the case, and why these results are broadly consistent between species, with possibly significant within-species variability, remains to be investigated further as it was beyond the goal of this methodological work. We note that, possibly, these results are affected by our choice to alternate sampling at midnight and at noon and to limit the length of trajectories to 500 locations, restricting the receptive field of our model to about 250 days. This may have weakened or excluded the influence of migration, which commonly leads to seasonal back-and-forth movement patterns. Therefore, in future work, it would be interesting to study how the temporal scale and resolution of trajectories affects the results. It is possible that using data with a higher temporal resolution and/or longer temporal context would reveal a second peak in context length, indicative of nested scale-patterns in the trajectories.

The effect of *spatial* scale and resolution of the geospatial variables may also be worth investigating. Instead of only considering the features at a given location, future work should explore including information from a local neighborhood. Apart from improving predictions in general, this additional context information may enable dealing with lower-accuracy (e.g. Argos) tracking data, where the variables associated with exact location may become too noisy.

One obvious limitation of our approach is the data requirement. As with all deep learning approaches, learning is limited by the data available in the training set, and enough data should also be available for validation and test sets. The whole dataset we gathered here, despite being rather large (*>* 1500 trajectories) compared to movement datasets currently analyzed in ecology, is likely close to the minimal size required to obtain a robust model and avoid severe overfitting issues. Currently, there are probably very few, if any, single-species datasets large enough to fit this model. For this reason, we aggregated data from numerous species; as a benefit, this allowed us to demonstrate that comparative analyses could be conducted with the model, for instance by comparing the distribution of relevant context lengths between species or higher-order taxa.

An important characteristic of the proposed approach is that the model not only accounts for past movements to predict new ones, but can also account for environmental predictors. First, this is crucial for realistic predictions, as the step-selection literature has well demonstrated that step selection by animals is critically linked to habitats to be traversed or reached. Second, this allows to evaluate the relative importance of predictors in improving predictions. Interestingly, we found that purely relative positional information (movement vectors) could be more important than environmental variables for future location prediction. We tentatively suggest that this result might be linked to the fact that most animals favor familiar places and by doing so restrict themselves to well-established home-ranges [180]. We however also found, without surprise, that among the environmental variables tested, land cover and human footprint significantly affected animal movements [181].

To summarize, in the present work, we provide a new, state-of-the art model to analyze and predict animal movement data. The novelty of the model lies in the fact that it leverages the power of deep learning approaches and can account for past movements in the predictions. However, we emphasize, and have shown above, that the model is not only a tool for prediction, but can also be used to test hypotheses about the intrinsic and extrinsic drivers of animal movements.

## 7. Data, script and code availability

Our code used for data retrieval, data pre-processing, model training, and analyses is available online as open source software^12^ [182]; the code for efficient geospatial variable loading is released as a separate open source package, gps2var^13^ [183]. We also release the weights of the trained models [184].

The data used in this work was compiled from 102 tracking studies, out of which:

- 98 are publicly available and were retrieved on 15 February 2022 via the Movebank API;
- more are additionally made publicly available in Movebank [166, 167] at the time of writing;
- the remaining 2 (plains zebras and blue wildebeest in Hluhluwe-iMfolozi Park) are under restricted use imposed by the institution managing the study area.

For a list of the Movebank studies, see Table 4.

## 8. Funding and acknowledgments

This work was supported by the LabEx NUMEV (ANR-10-LABX-0020) and the REPOS project, both funded by the I-Site MUSE (ANR-16-IDEX-0006). Computations were performed using HPC/AI resources from GENCI-IDRIS (Grant AD011012019R1).

We would like to thank all authors who made their data available through Movebank under Creative Commons licenses.

## 9. Conflict of interest disclosure

The authors declare they have no conflict of interest relating to the content of this article.

## Supplementary information

**Table 4:**
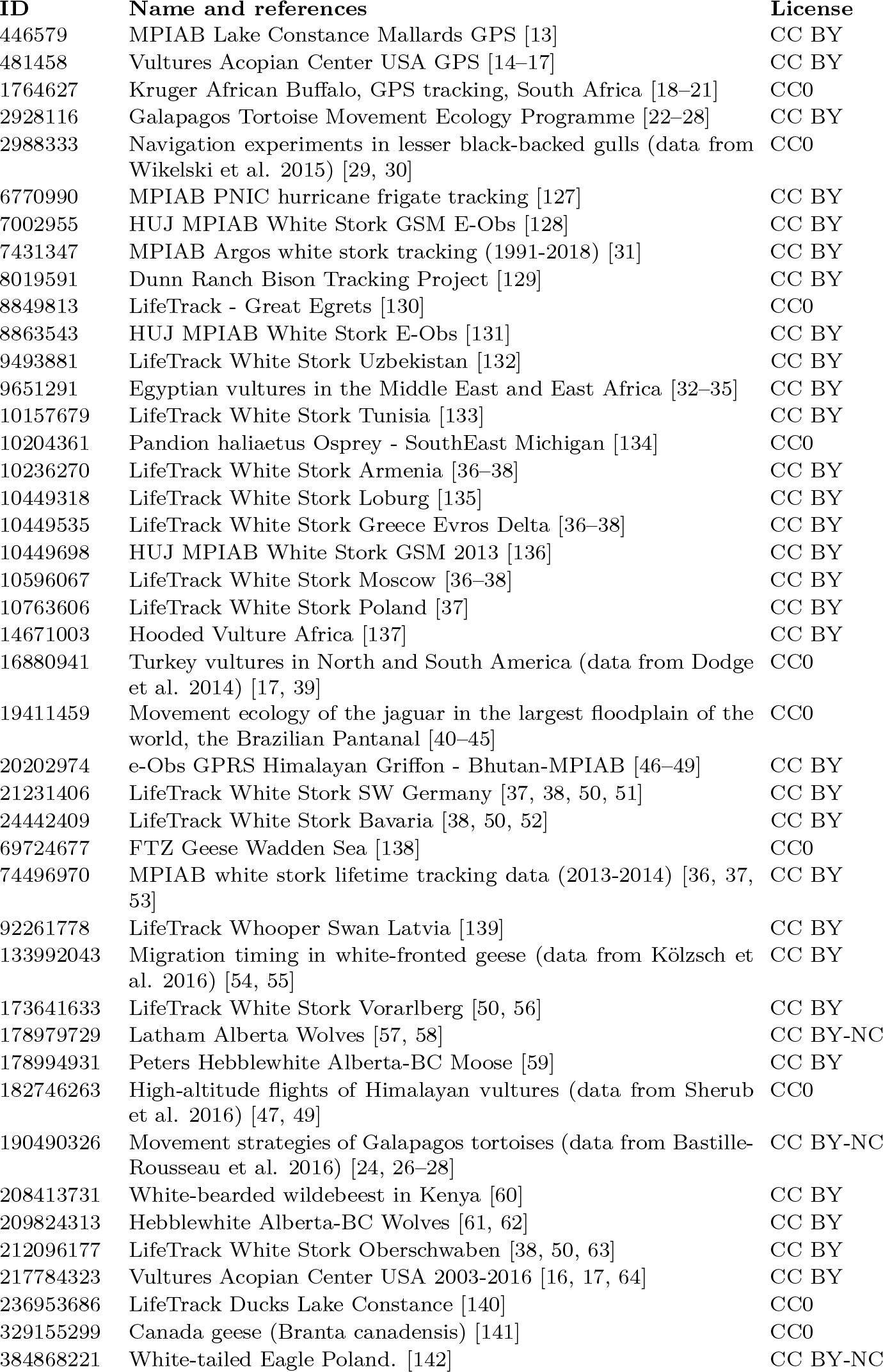
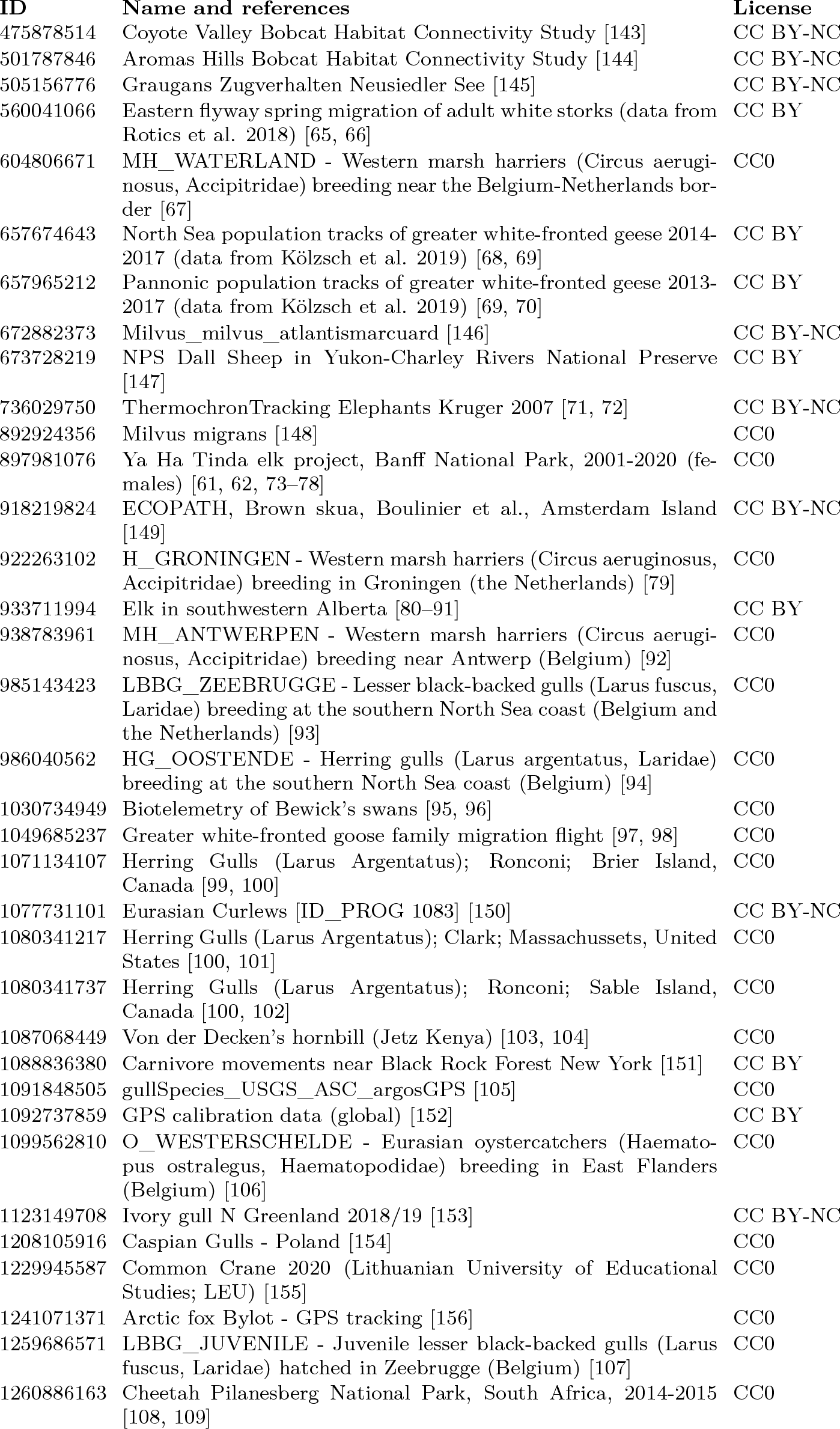
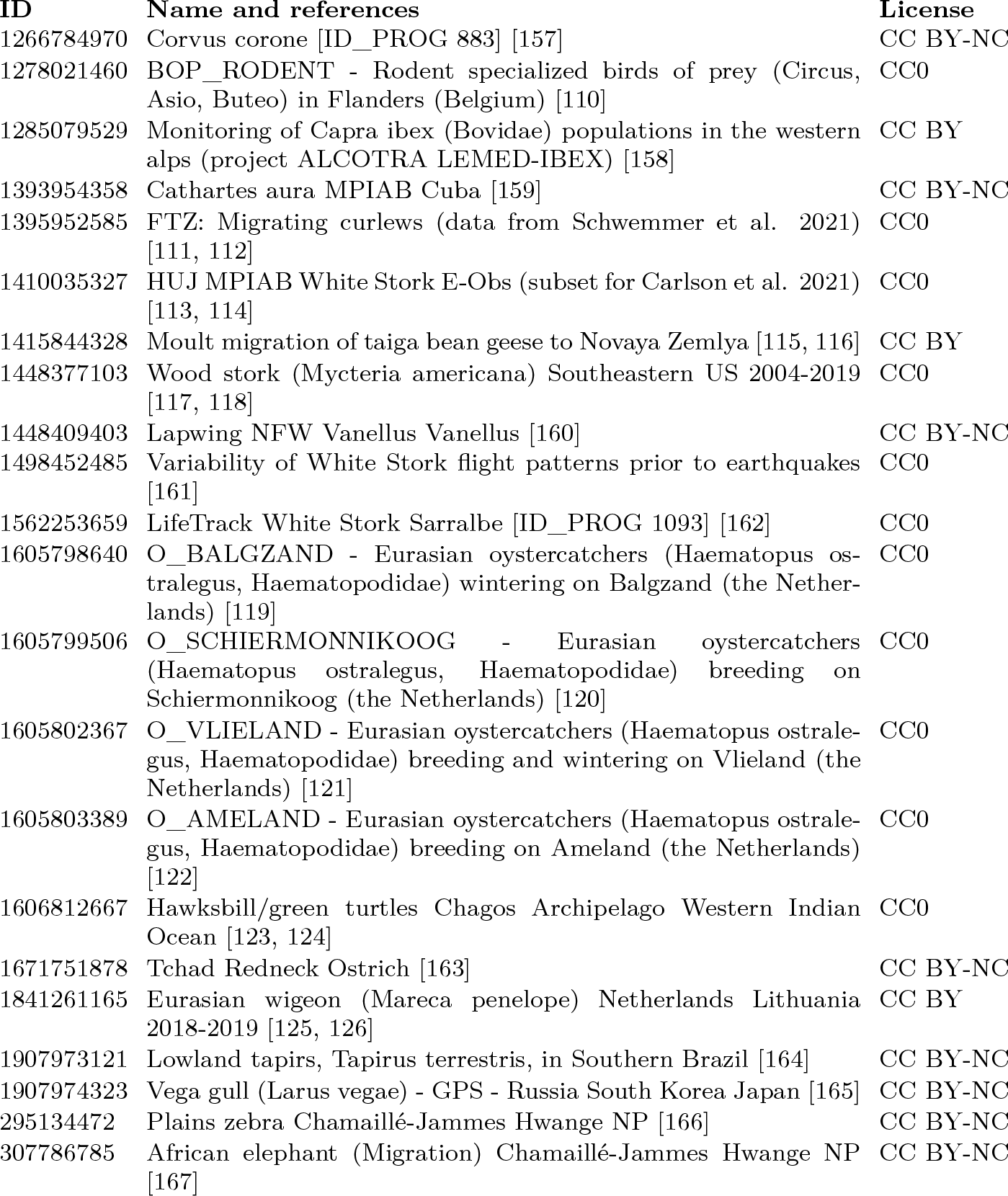
The list of all Movebank datasets used in this work.

**Table 5:**
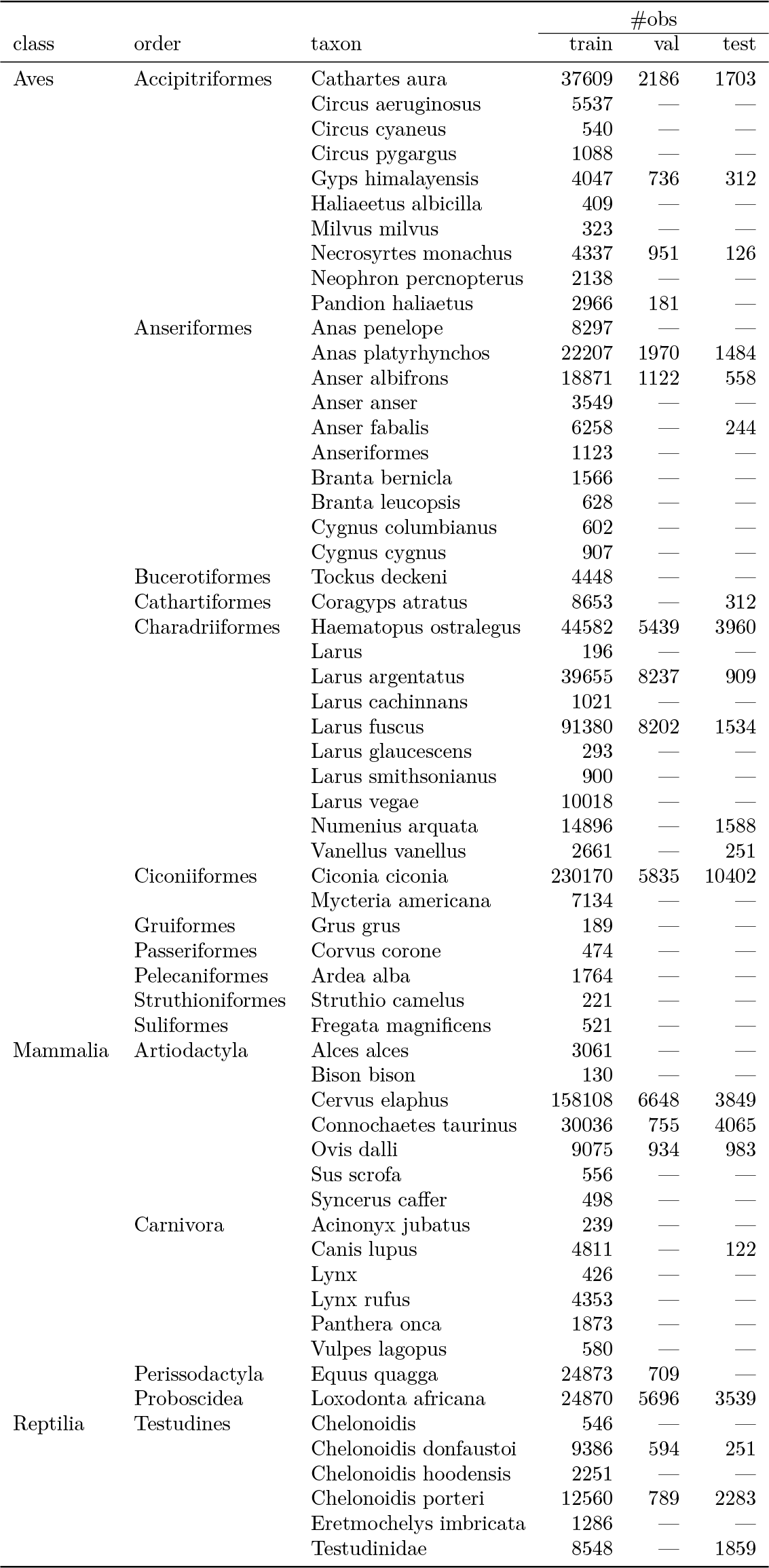
Number of observations of each taxon in each section of the dataset.

**Table 6:** WorldClim bioclimatic variables as listed at https://www.worldclim.org/data/bioclim.html.

| Variable | Description |
| --- | --- |
| BIO1 | Annual Mean Temperature |
| BIO2 | Mean Diurnal Range (mean of monthly (max temp – min temp)) |
| BIO3 | Isothermality ($\text{BIO2}/\text{BIO7} \times 100$) |
| BIO4 | Temperature Seasonality (standard deviation $\times 100$) |
| BIO5 | Max Temperature of Warmest Month |
| BIO6 | Min Temperature of Coldest Month |
| BIO7 | Temperature Annual Range ($\text{BIO5} - \text{BIO6}$) |
| BIO8 | Mean Temperature of Wettest Quarter |
| BIO9 | Mean Temperature of Driest Quarter |
| BIO10 | Mean Temperature of Warmest Quarter |
| BIO11 | Mean Temperature of Coldest Quarter |
| BIO12 | Annual Precipitation |
| BIO13 | Precipitation of Wettest Month |
| BIO14 | Precipitation of Driest Month |
| BIO15 | Precipitation Seasonality (Coefficient of Variation) |
| BIO16 | Precipitation of Wettest Quarter |
| BIO17 | Precipitation of Driest Quarter |
| BIO18 | Precipitation of Warmest Quarter |
| BIO19 | Precipitation of Coldest Quarter |

## Footnotes

1 https://github.com/cifkao/moveformer

2 www.movebank.org

3 as of 15 February 2022

4 https://creativecommons.org/

5 Specifically, after filtering out infrequent species (with less than 10 members in the full dataset), we randomly assign 4 % of the remaining individuals to the validation set, another 5 % to the test set, and the rest to the training set.

6 A Transformer typically supports sequences up to a maximum length, determined by the lengths of the sequences encountered during training. Training on full-length sequences is usually not feasible due to its computational cost, but also because the training dataset is likely to only contain a *single* sequence of exactly the full length, preventing successful generalization up to that length. Setting an upper length limit close to the median trajectory length is a convenient way to avoid these issues.

7 We use *ξ*_1…*N*_ as shorthand notation for the sequence *ξ*_1_, *ξ*_2_, …, *ξ*_*N*_ . Note that *N* may be different for each trajectory in the dataset.

8 https://pytorch.org/

9 https://github.com/lucidrains/x-transformers

10 https://github.com/rasterio/rasterio

11 We use the number of candidates *K* as the base of the logarithm for consistency with Eq. (5) and noting that this amounts to a multiplicative constant (1*/* log *K*).

12 https://github.com/cifkao/moveformer

13 https://github.com/cifkao/gps2var

## Notes

https://github.com/cifkao/moveformer

https://doi.org/10.5281/zenodo.7698263

https://github.com/cifkao/gps2var

https://doi.org/10.5281/zenodo.8217156

### Competing Interest Statement

The authors have declared no competing interest.

### Summary of Updates

Added PCIEcology recommendation badge.

## References

[1] Roland Kays, Margaret C. Crofoot, Walter Jetz, and Martin Wikelski. Terrestrial animal tracking as an eye on life and planet. Science, 348(6240): aaa2478, 2015. Publisher: American Association for the Advancement of Science.

[2] Henrik Thurfjell, Simone Ciuti, and Mark S. Boyce. Applications of step-selection functions in ecology and conservation. Movement ecology, 2:1–12, 2014. Publisher: Springer.

[3] Tal Avgar, Jonathan R. Potts, Mark A. Lewis, and Mark S. Boyce. Integrated step selection analysis: bridging the gap between resource selection and animal movement. Methods in Ecology and Evolution, 7(5):619–630, 2016. ISSN 2041-210X. doi: 10.1111/2041-210X.12528. URL https://onlinelibrary.wiley.com/doi/abs/10.1111/2041-210X.12528.

[4] John Fieberg, Johannes Signer, Brian Smith, and Tal Avgar. A ‘How to’ guide for interpreting parameters in habitat-selection analyses. Journal of Animal Ecology, 5(5):1027–1043, 2021. ISSN 1365-2656. doi: 10.1111/1365-2656.13441. URL https://onlinelibrary.wiley.com/doi/abs/10.1111/1365-2656.13441.

[5] William F. Fagan, Mark A. Lewis, Marie Auger-Méthé, Tal Avgar, Simon Benhamou, Greg Breed, Lara LaDage, Ulrike E. Schlägel, Wen-wu Tang, and Yannis P. Papastamatiou. Spatial memory and animal movement. Ecology letters, 10(10):1316–1329, 2013. Publisher: Wiley Online Library.

[6] Louise Riotte-Lambert, Simon Benhamou, and Simon Chamaillé-Jammes. How memory-based movement leads to nonterritorial spatial segregation. The American Naturalist, 4(4):E103–E116, 2015. Publisher: University of Chicago Press Chicago, IL.

[7] Nathan Ranc, Paul R. Moorcroft, Federico Ossi, and Francesca Cagnacci. Experimental evidence of memory-based foraging decisions in a large wild mammal. Proceedings of the National Academy of Sciences, 118(15): e2014856118, 2021. Publisher: National Acad Sciences.

[8] Luiz Gustavo R. Oliveira-Santos, James D. Forester, Ubiratan Piovezan, Walfrido M. Tomas, and Fernando A. S. Fernandez. Incorporating animal spatial memory in step selection functions. Journal of Animal Ecology, 2(2):516–524, 2016. ISSN 1365-2656. doi: 10.1111/1365-2656.12485. URL https://onlinelibrary.wiley.com/doi/abs/10.1111/1365-2656.12485. _eprint: https://besjournals.onlinelibrary.wiley.com/doi/pdf/10.1111/1365-2656.12485.

[9] Jerod A. Merkle, Hall Sawyer, Kevin L. Monteith, Samantha PH Dwinnell, Gary L. Fralick, and Matthew J. Kauffman. Spatial memory shapes migration and its benefits: evidence from a large herbivore. Ecology letters, 11(11):1797–1805, 2019. Publisher: Wiley Online Library.

[10] Oded Berger-Tal and Shirli Bar-David. Recursive movement patterns: review and synthesis across species. Ecosphere, 9(9):art149, 2015. ISSN 2150-8925. doi: 10.1890/ES15-00106.1. URL https://onlinelibrary.wiley.com/doi/abs/10.1890/ES15-00106.1. _eprint: https://esajournals.onlinelibrary.wiley.com/doi/pdf/10.1890/ES15-00106.1.

[11] Louise Riotte-Lambert, Simon Benhamou, and Simon Chamaillé-Jammes. From randomness to traplining: a framework for the study of routine movement behavior. Behavioral Ecology, 1(1):280–287, January 2017. ISSN 1045-2249. doi: 10.1093/beheco/arw154. URL 10.1093/beheco/arw154.

[12] Roland Kays, Sarah C. Davidson, Matthias Berger, Gil Bohrer, Wolfgang Fiedler, Andrea Flack, Julian Hirt, Clemens Hahn, Dominik Gauggel, Benedict Russell, Andrea Kölzsch, Ashley Lohr, Jesko Partecke, Michael Quetting, Kamran Safi, Anne Scharf, Gabriel Schneider, Ilona Lang, Friedrich Schaeuffelhut, Matthias Landwehr, Martin Storhas, Louis van Schalkwyk, Candace Vinciguerra, Rolf Weinzierl, and Martin Wikelski. The Movebank system for studying global animal movement and demography. Methods in Ecology and Evolution, 2(2):419–431, 2022. ISSN 2041-210X. doi: 10.1111/2041-210X.13767. URL 10.1111/2041-210X.13767. https://onlinelibrary.wiley.com/doi/pdf/10.1111/2041-210X.13767.

[13] Pius Korner, Annette Sauter, Wolfgang Fiedler, and Lukas Jenni. Variable allocation of activity to daylight and night in the mallard. Animal Behaviour, 115:69–79, may 2016. doi: 10.1016/j.anbehav.2016.02.026. URL 10.1016/j.anbehav.2016.02.026.

[14] Keith L. Bildstein, David Barber, Marc J. Bechard, Maricel Graña Grilli, and Jean-François Therrien. Data from: Study “Vultures Acopian Center USA GPS” (2003-2021), 2021. URL 10.5441/001/1.f3qt46r2.

[15] Julie M. Mallon, Keith L. Bildstein, and William F. Fagan. Inclement weather forces stopovers and prevents migratory progress for obligate soaring migrants. Movement Ecology, 9(1), jul 2021. doi: 10.1186/s40462-021-00274-6. URL 10.1186/s40462-021-00274-6.

[16] Maricel Graña Grilli, Sergio A. Lambertucci, Jean-François Therrien, and Keith L. Bildstein. Wing size but not wing shape is related to migratory behavior in a soaring bird. Journal of Avian Biology, 5(5):669–678, mar 2017. doi: 10.1111/jav.01220. URL 10.1111/jav.01220.

[17] Somayeh Dodge, Gil Bohrer, Keith Bildstein, Sarah C. Davidson, Rolf Weinzierl, Marc J. Bechard, David Barber, Roland Kays, David Brandes, Jiawei Han, and Martin Wikelski. Environmental drivers of variability in the movement ecology of turkey vultures (Cathartes aura ) in North and South America. Philosophical Transactions of the Royal Society B: Biological Sciences, 1643(1643):20130195, may 2014. doi: 10.1098/rstb.2013.0195. URL 10.1098/rstb.2013.0195.

[18] Paul C. Cross, Justin A. Bowers, Craig T. Hay, Julie Wolhuter, Peter Buss, Markus Hofmeyr, Johan T. Du Toit, and Wayne M. Getz. Data from: Nonparameteric kernel methods for constructing home ranges and utilization distributions, 2016. URL 10.5441/001/1.j900f88t.

[19] Justin M. Calabrese, Chris H. Fleming, and Eliezer Gurarie. ctmm: an r package for analyzing animal relocation data as a continuous-time stochastic process. Methods in Ecology and Evolution, 9(9):1124–1132, may 2016. doi: 10.1111/2041-210x.12559. URL 10.1111/2041-210X.12559.

[20] P. C. Cross, D. M. Heisey, J. A. Bowers, C. T. Hay, J. Wolhuter, P. Buss, M. Hofmeyr, A. L. Michel, R. G. Bengis, T. L. F. Bird, J. T. Du Toit, and W. M. Getz. Disease, predation and demography: assessing the impacts of bovine tuberculosis on African buffalo by monitoring at individual and population levels. Journal of Applied Ecology, 2(2):467–475, apr 2009. doi: 10.1111/j.1365-2664.2008.01589.x. URL 10.1111/j.1365-2664.2008.01589.x.

[21] Wayne M. Getz, Scott Fortmann-Roe, Paul C. Cross, Andrew J. Lyons, Sadie J. Ryan, and Christopher C. Wilmers. LoCoH: Nonparameteric kernel methods for constructing home ranges and utilization distributions. PLoS ONE, 2(2):e207, feb 2007. doi: 10.1371/journal.pone.0000207. URL 10.1371/journal.pone.0000207.

[22] Guillaume Bastille-Rousseau, Charles B. Yackulic, James Gibbs, Jacqueline L. Frair, Freddy Cabrera, and Stephen Blake. Data from: Migration triggers in a large herbivore: Galápagos giant tortoises navigating resource gradients on volcanoes, 2019. URL 10.5441/001/1.6gr485fk.

[23] Guillaume Bastille-Rousseau, Charles B. Yackulic, Jacqueline L. Frair, Freddy Cabrera, and Stephen Blake. Data from: Allometric and temporal scaling of movement characteristics in Galapagos tortoises, 2016. URL 10.5441/001/1.2cp86266.

[24] Guillaume Bastille-Rousseau, Jonathan R. Potts, Charles B. Yackulic, Jacqueline L. Frair, E. Hance Ellington, and Stephen Blake. Data from: Flexible characterization of animal movement pattern using net squared displacement and a latent state model, 2016. URL 10.5441/001/1.356nb5mf.

[25] Guillaume Bastille-Rousseau, Charles B. Yackulic, James P. Gibbs, Jacqueline L. Frair, Freddy Cabrera, and Stephen Blake. Migration triggers in a large herbivore: Galápagos giant tortoises navigating resource gradients on volcanoes. Ecology, 6(6):e02658, apr 2019. doi: 10.1002/ecy.2658. URL 10.1002/ecy.2658.

[26] Guillaume Bastille-Rousseau, James P. Gibbs, Charles B. Yackulic, Jacqueline L. Frair, Fredy Cabrera, Louis-Philippe Rousseau, Martin Wikelski, Franz Kümmeth, and Stephen Blake. Animal movement in the absence of predation: environmental drivers of movement strategies in a partial migration system. Oikos, 7(7):1004–1019, jan 2017. doi: 10.1111/oik.03928. URL 10.1111/oik.03928.

[27] Guillaume Bastille-Rousseau, Jonathan R. Potts, Charles B. Yackulic, Jacqueline L. Frair, E. Hance Ellington, and Stephen Blake. Flexible characterization of animal movement pattern using net squared displacement and a latent state model. Movement Ecology, 4(1), jun 2016. doi: 10.1186/s40462-016-0080-y. URL 10.1186/s40462-016-0080-y.

[28] Guillaume Bastille-Rousseau, Charles B. Yackulic, Jacqueline L. Frair, Freddy Cabrera, and Stephen Blake. Allometric and temporal scaling of movement characteristics in Galapagos tortoises. Journal of Animal Ecology, 5(5):1171–1181, jul 2016. doi: 10.1111/1365-2656.12561. URL 10.1111/1365-2656.12561.

[29] Martin Wikelski, Elena Arriero, Anna Gagliardo, Richard Holland, Markku J. Huttunen, Risto Juvaste, Inge Mueller, Grigori Tertitski, Kasper Thorup, Martin Wild, Markku Alanko, Franz Bairlein, Alexander Cherenkov, Alison Cameron, Reinhard Flatz, Juhani Hannila, Ommo Hüppop, Markku Kangasniemi, Bart Kranstauber, Maija-Liisa Penttinen, Kamran Safi, Vladimir Semashko, Heidi Schmid, and Ralf Wistbacka. Data from: True navigation in migrating gulls requires intact olfactory nerves, 2015. URL 10.5441/001/1.q986rc29.

[30] Martin Wikelski, Elena Arriero, Anna Gagliardo, Richard A. Holland, Markku J. Huttunen, Risto Juvaste, Inge Mueller, Grigori Tertitski, Kasper Thorup, Martin Wild, Markku Alanko, Franz Bairlein, Alexander Cherenkov, Alison Cameron, Reinhard Flatz, Juhani Hannila, Ommo Hüppop, Markku Kangasniemi, Bart Kranstauber, Maija-Liisa Penttinen, Kamran Safi, Vladimir Semashko, Heidi Schmid, and Ralf Wistbacka. True navigation in migrating gulls requires intact olfactory nerves. Scientific Reports, 5(1), nov 2015. doi: 10.1038/srep17061. URL 10.1038/srep17061.

[31] Andrea Kölzsch, Erik Kleyheeg, Helmut Kruckenberg, Michael Kaatz, and Bernd Blasius. A periodic Markov model to formalize animal migration on a network. Royal Society Open Science, 6(6):180438, jun 2018. doi: 10.1098/rsos.180438. URL 10.1098/rsos.180438.

[32] Evan R. Buechley and Çağan H. Şekercioğlu. Data from: Satellite tracking a wide-ranging endangered vulture species to target conservation actions in the Middle East and East Africa, 2019. URL 10.5441/001/1.385gk270.

[33] W. Louis Phipps, Pascual López-López, Evan R. Buechley, Steffen Oppel, Ernesto Álvarez, Volen Arkumarev, Rinur Bekmansurov, Oded Berger-Tal, Ana Bermejo, Anastasios Bounas, Isidoro Carbonell Alanís, Javier de la Puente, Vladimir Dobrev, Olivier Duriez, Ron Efrat, Guillaume Fréchet, Javier García, Manuel Galán, Clara García-Ripollés, Alberto Gil, Juan José Iglesias-Lebrija, José Jambas, Igor V. Karyakin, Erick Kobierzycki, Elzbieta Kret, Franziska Loercher, Antonio Monteiro, Jon Morant Etxebarria, Stoyan C. Nikolov, José Pereira, Lubomír Peške, Cecile Ponchon, Eduardo Realinho, Victoria Saravia, Cağan H. Sekercioğlu, Theodora Skartsi, José Tavares, Joaquim Teodósio, Vicente Urios, and Núria Vallverdú. Spatial and Temporal Variability in Migration of a Soaring Raptor Across Three Continents. Frontiers in Ecology and Evolution, 7, sep 2019. doi: 10.3389/fevo.2019.00323. URL 10.3389/fevo.2019.00323.

[34] Evan R. Buechley, Michael J. McGrady, Emrah Çoban, and Çağan H. Şekercioğlu. Satellite tracking a wide-ranging endangered vulture species to target conservation actions in the Middle East and East Africa. Biodiversity and Conservation, 9(9):2293–2310, mar 2018. doi: 10.1007/s10531-018-1538-6. URL 10.1007/s10531-018-1538-6.

[35] Evan R. Buechley, Steffen Oppel, William S. Beatty, Stoyan C. Nikolov, Vladimir Dobrev, Volen Arkumarev, Victoria Saravia, Clementine Bougain, Anastasios Bounas, Elzbieta Kret, Theodora Skartsi, Lale Aktay, Karen Aghababyan, Ethan Frehner, and Çağan H. Şekercioğlu. Identifying critical migratory bottlenecks and high-use areas for an endangered migratory soaring bird across three continents. Journal of Avian Biology, 7(7):e01629, jul 2018. doi: 10.1111/jav.01629. URL 10.1111/jav.01629.

[36] Andrea Flack, Wolfgang Fiedler, Julio Blas, Ivan Pokrovski, B. Mitropolsky, Michael Kaatz, Karen Aghababyan, A. Khachatryan, Ioannis Fakriadis, Eleni Makrigianni, Leszek Jerzak, M. Shamin, C. Shamina, H. Azafzaf, Claudia Feltrup-Azafzaf, Thabiso M. Mokotjomela, and Martin Wikelski. Data from: Costs of migratory decisions: a comparison across eight white stork populations, 2015. URL 10.5441/001/1.78152p3q.

[37] Andrea Flack, Wolfgang Fiedler, Julio Blas, Ivan Pokrovsky, Michael Kaatz, Maxim Mitropolsky, Karen Aghababyan, Ioannis Fakriadis, Eleni Makrigianni, Leszek Jerzak, Hichem Azafzaf, Claudia Feltrup-Azafzaf, Shay Rotics, Thabiso M. Mokotjomela, Ran Nathan, and Martin Wikelski. Costs of migratory decisions: A comparison across eight white stork populations. Science Advances, 2(1), jan 2016. doi: 10.1126/sciadv.1500931. URL 10.1126/sciadv.1500931.

[38] Roland Kays, Sarah C. Davidson, Matthias Berger, Gil Bohrer, Wolfgang Fiedler, Andrea Flack, Julian Hirt, Clemens Hahn, Dominik Gauggel, Benedict Russell, Andrea Kölzsch, Ashley Lohr, Jesko Partecke, Michael Quetting, Kamran Safi, Anne Scharf, Gabriel Schneider, Ilona Lang, Friedrich Schaeuffelhut, Matthias Landwehr, Martin Storhas, Louis van Schalkwyk, Candace Vinciguerra, Rolf Weinzierl, and Martin Wikelski. The Movebank system for studying global animal movement and demography. Methods in Ecology and Evolution, 2(2):419–431, ec 2021. doi: 10.1111/2041-210x.13767. URL 10.1111/2041-210X.13767.

[39] Keith Bildstein, David Barber, and Marc J. Bechard. Data from: Environmental drivers of variability in the movement ecology of turkey vultures (Cathartes aura) in North and South America, 2014. URL 10.5441/001/1.46ft1k05.

[40] Ronaldo Gonçalves Morato, Daniel Luiz Zanella Kantek, Samiko Miyazaki, Thadeu Deluque, and Rogerio Cunha De Paula. Data from: Jaguar movement database—a GPS-based movement dataset of an apex predator in the Neotropics, 2021. URL 10.5441/001/1.3c4fv0m4.

[41] Ronaldo G. Morato, Jeffrey J. Thompson, Agustin Paviolo, Jesus A. de La Torre, Fernando Lima, Roy T. McBride, Rogerio C. Paula, Laury Cullen, Leandro Silveira, Daniel L. Z. Kantek, Emiliano E. Ramalho, Louise Maranhão, Mario Haberfeld, Denis A. Sana, Rodrigo A. Medellin, Eduardo Carrillo, Victor Montalvo, Octavio Monroy-Vilchis, Paula Cruz, Anah T. Jacomo, Natalia M. Torres, Giselle B. Alves, Ivonne Cassaigne, Ron Thompson, Carolina Saens-Bolanos, Juan Carlos Cruz, Luiz D. Alfaro, Isabel Hagnauer, Xavier Marina da Silva, Alexandre Vogliotti, Marcela F. D. Moraes, Selma S. Miyazaki, Thadeu D. C. Pereira, Gediendson R. Araujo, Leanes Cruz da Silva, Lucas Leuzinger, Marina M. Carvalho, Lilian Rampin, Leonardo Sartorello, Howard Quigley, Fernando Tortato, Rafael Hoogesteijn, Peter G. Crawshaw, Allison L. Devlin, Joares A. May, Fernando C. C. de Azevedo, Henrique V. B. Concone, Veronica A. Quiroga, Sebastian A. Costa, Juan P. Arrabal, Ezequiel Vanderhoeven, Yamil E. Di Blanco, Alexandre M. C. Lopes, Cynthia E. Widmer, and Milton Cezar Ribeiro. Jaguar movement database: a GPS-based movement dataset of an apex predator in the neotropics. Ecology, 7(7):1691–1691, jul 2018. doi: 10.1002/ecy.2379. URL 10.1002/ecy.2379.

[42] R.G. Morato, G.M. Connette, J.A. Stabach, R.C. De Paula, K.M.P.M. Ferraz, D.L.Z. Kantek, S.S. Miyazaki, T.D.C. Pereira, L.C. Silva, A. Paviolo, C. De Angelo, M.S. Di Bitetti, P. Cruz, F. Lima, L. Cullen, D.A. Sana, E.E. Ramalho, M.M. Carvalho, M.X. da Silva, M.D.F. Moraes, A. Vogliotti, J.A. May, M. Haberfeld, L. Rampim, L. Sartorello, G.R. Araujo, G. Wittemyer, M.C. Ribeiro, and P. Leimgruber. Resource selection in an apex predator and variation in response to local landscape characteristics. Biological Conservation, 228:233–240, ec 2018. doi: 10.1016/j.biocon.2018.10.022. URL 10.1016/j.biocon.2018.10.022.

[43] Ronaldo G. Morato, Jared A. Stabach, Chris H. Fleming, Justin M. Calabrese, Rogério C. De Paula, Kátia M. P. M. Ferraz, Daniel L. Z. Kantek, Selma S. Miyazaki, Thadeu D. C. Pereira, Gediendson R. Araujo, Agustin Paviolo, Carlos De Angelo, Mario S. Di Bitetti, Paula Cruz, Fernando Lima, Laury Cullen, Denis A. Sana, Emiliano E. Ramalho, Marina M. Carvalho, Fábio H. S. Soares, Barbara Zimbres, Marina X. Silva, Marcela D. F. Moraes, Alexandre Vogliotti, Joares A. May, Mario Haberfeld, Lilian Rampim, Leonardo Sartorello, Milton C. Ribeiro, and Peter Leimgruber. Space Use and Movement of a Neotropical Top Predator: The Endangered Jaguar. PLOS ONE, 12(12):e0168176, ec 2016. doi: 10.1371/journal.pone.0168176. URL 10.1371/journal.pone.0168176.

[44] Ronaldo G. Morato, Jeffrey J. Thompson, Agustín Paviolo, J. Antonio De La Torre, Fernando Lima, McBride Jr., Roy T., Rogério C. Paula, Cullen Jr., Laury, Leandro Silveira, Daniel L.Z. Kantek, Emiliano E. Ramalho, Louise Maranhão, Mario Haberfeld, Denis A. Sana, Rodrigo A. Medellin, Eduardo Carrillo, Victor Montalvo, Octavio Monroy-Vilchis, Paula Cruz, Anah Tereza Jácomo, Natalia M. Torres, Giselle B. Alves, Ivonne Cassaigne, Ron Thompson, Carolina Saenz-Bolanos, Juan Carlos Cruz, Luis D. Alfaro, Isabel Hagnauer, Marina Xavier Da Silva, Alexandre Vogliotti, Marcela Figuerêdo Duarte Moraes, Selma S. Miyazaki, Thadeu D.C. Pereira, Gediendson R. Araujo, Leanes Cruz Da Silva, Lukas Leuzinger, Marina M Carvalho, Lilian Rampim, Leonardo Sartorello, Howard Quigley, Fernando Tortato, Rafael Hoogesteijn, Crawshaw Jr., Peter G., Allison L. Devlin, May Jr., Joares A., Fernando C.C. De Azevedo, Henrique Villas Boas Concone, Veronica A. Quiroga, Sebastián A. Costa, Juan P. Arrabal, Ezequiel Vanderhoeven, Yamil E. Di Blanco, Alexandre M.C. Lopes, Cynthia E. Widmer, Milton Cezar Ribeiro, Carolina Saens-Bolanos, Luiz D. Alfaro, and Joares A. May. Data from: Jaguar Movement Database: a GPS-based movement dataset of an apex predator in the Neotropics, 2019. URL 10.5061/dryad.2dh0223.

[45] Miltinhoastronauta. Leeclab/Jaguar_Movement: Jaguar Movement Database V1.0 Released!, 2018. URL 10.5281/zenodo.1345119.

[46] Sherub Sherub and Martin Wikelski. Data from: Longer days enable higher diurnal activity for migratory birds [Himalayan griffons], 2021. URL 10.5441/001/1.4n2501f5.

[47] Sherub Sherub, Martin Wikelski, Wolfgang Fiedler, and Sarah C. Davidson. Data from: Behavioural adaptations to flight into thin air, 2016. URL 10.5441/001/1.143v2p2k.

[48] Ivan Pokrovsky, Andrea Kölzsch, Sherub Sherub, Wolfgang Fiedler, Peter Glazov, Olga Kulikova, Martin Wikelski, and Andrea Flack. Longer days enable higher diurnal activity for migratory birds. Journal of Animal Ecology, 9(9):2161–2171, mar 2021. doi: 10.1111/1365-2656.13484. URL 10.1111/1365-2656.13484.

[49] Sherub Sherub, Gil Bohrer, Martin Wikelski, and Rolf Weinzierl. Behavioural adaptations to flight into thin air. Biology Letters, 12(10): 20160432, oct 2016. doi: 10.1098/rsbl.2016.0432. URL 10.1098/rsbl.2016.0432.

[50] Yachang Cheng, Wolfgang Fiedler, Martin Wikelski, and Andrea Flack. “closer-to-home” strategy benefits juvenile survival in a long-distance migratory bird. Ecology and Evolution, 16(16):8945–8952, jul 2019. doi: 10.1002/ece3.5395. URL 10.1002/ece3.5395.

[51] Rolf Weinzierl, Gil Bohrer, Bart Kranstauber, Wolfgang Fiedler, Martin Wikelski, and Andrea Flack. Wind estimation based on thermal soaring of birds. Ecology and Evolution, 24(24):8706–8718, nov 2016. doi: 10.1002/ece3.2585. URL 10.1002/ece3.2585.

[52] Wolfgang Fiedler, Elke Leppelsack, Hans Leppelsack, Thomas Stahl, Oda Wieding, and Martin Wikelski. Data from: Study “LifeTrack White Stork Bavaria” (2014-2019), 2019. URL 10.5441/001/1.v1cs4nn0.

[53] Roland Kays, Margaret C. Crofoot, Walter Jetz, and Martin Wikelski. Terrestrial animal tracking as an eye on life and planet. Science, 348 (6240), jun 2015. doi: 10.1126/science.aaa2478. URL 10.1126/science.aaa2478.

[54] Andrea Kölzsch, Helmut Kruckenberg, Peter Glazov, Gerhard J.D.M. Müskens, and Martin Wikelski. Data from: Towards a new understanding of migration timing: slower spring than autumn migration in geese reflects different decision rules for stopover use and departure, 2016. URL 10.5441/001/1.31c2v92f.

[55] Andrea Kölzsch, Gerhard J. D. M. Müskens, Helmut Kruckenberg, Peter Glazov, Rolf Weinzierl, Bart A. Nolet, and Martin Wikelski. Towards a new understanding of migration timing: slower spring than autumn migration in geese reflects different decision rules for stopover use and departure. Oikos, 10(10):1496–1507, feb 2016. doi: 10.1111/oik.03121. URL 10.1111/oik.03121.

[56] Wolfgang Fiedler, Walter Niederer, Alwin Schönenberger, Andrea Flack, and Martin Wikelski. Data from: Study “LifeTrack White Stork Vorarlberg” (2016-2019), 2019. URL 10.5441/001/1.71r7pp6q.

[57] A. David M. Latham and Stan Boutin. Data from: Wolf ecology and caribou-primary prey-wolf spatial relationships in low productivity peatland complexes in northeastern Alberta, 2019. URL 10.5441/001/1.7vr1k987.

[58] A. David M. Latham, M. Cecilia Latham, Mark S. Boyce, and Stan Boutin. Movement responses by wolves to industrial linear features and their effect on woodland caribou in northeastern Alberta. Ecological Applications, 8(8):2854–2865, ec 2011. doi: 10.1890/11-0666.1. URL 10.1890/11-0666.1.

[59] Wibke Peters, Mark Hebblewhite, Nicholas DeCesare, Francesca Cagnacci, and Marco Musiani. Resource separation analysis with moose indicates threats to caribou in human altered landscapes. Ecography, 4(4):487–498, oct 2012. doi: 10.1111/j.1600-0587.2012.07733.x. URL 10.1111/j.1600-0587.2012.07733.x.

[60] Jared A. Stabach, Lacey F. Hughey, Robin S. Reid, Jeffrey S. Worden, Peter Leimgruber, and Randall B. Boone. Data from: Study “White-bearded wildebeest in Kenya”, 2020. URL 10.5441/001/1.h0t27719.

[61] Mark Hebblewhite and Evelyn Merrill. Modelling wildlife-human relationships for social species with mixed-effects resource selection models. Journal of Applied Ecology, 3(3):834–844, jul 2007. doi: 10.1111/j.1365-2664.2008.01466.x. URL 10.1111/j.1365-2664.2008.01466.x.

[62] Mark Hebblewhite and Evelyn H. Merrill. Multiscale wolf predation risk for elk: does migration reduce risk? Oecologia, 2(2):377–387, feb 2007. doi: 10.1007/s00442-007-0661-y. URL 10.1007/s00442-007-0661-y.

[63] Wolfgang Fiedler, Andrea Flack, Andreas Schmidt, Ute Reinhard, and Martin Wikelski. Data from: Study “LifeTrack White Stork Oberschwaben” (2014-2019), 2019. URL 10.5441/001/1.c42j3js7.

[64] Keith L. Bildstein, David Barber, Marc J. Bechard, and Maricel Graña Grilli. Data from: Wing size but not wing shape is related to migratory behavior in a soaring bird, 2016. URL 10.5441/001/1.37r2b884.

[65] Shay Rotics, Michael Kaatz, Sondra Turjeman, Damaris Zurell, Martin Wikelski, Nir Sapir, Ute Eggers, Wolfgang Fiedler, Florian Jeltsch, and Ran Nathan. Data from: Early arrival at breeding grounds: causes, costs and a trade-off with overwintering latitude, 2018. URL 10.5441/001/1.v8d24552.

[66] Shay Rotics, Michael Kaatz, Sondra Turjeman, Damaris Zurell, Martin Wikelski, Nir Sapir, Ute Eggers, Wolfgang Fiedler, Florian Jeltsch, and Ran Nathan. Early arrival at breeding grounds: Causes, costs and a trade-off with overwintering latitude. Journal of Animal Ecology, 87(6): 1627–1638, oct 2018. doi: 10.1111/1365-2656.12898. URL 10.1111/1365-2656.12898.

[67] Anny Anselin, Peter Desmet, Tanja Milotic, Kjell Janssens, Filiep T’Jollyn, Luc De Bruyn, and Willem Bouten. MH_WATERLAND - Western marsh harriers (Circus aeruginosus, Accipitridae) breeding near the Belgium-Netherlands border, 2022. URL 10.5281/zenodo.3532940.

[68] Andrea Kölzsch, Gerhard J.D.M. Müskens, Sander Moonen, Helmut Kruckenberg, Peter Glazov, and Martin Wikelski. Data from: Flyway connectivity and exchange primarily driven by moult migration in geese [North Sea population], 2019. URL 10.5441/001/1.ct72m82n.

[69] A. Kölzsch, G. J. D. M. Müskens, P. Szinai, S. Moonen, P. Glazov, H. Kruckenberg, M. Wikelski, and B. A. Nolet. Flyway connectivity and exchange primarily driven by moult migration in geese. Movement Ecology, 7(1), jan 2019. doi: 10.1186/s40462-019-0148-6. URL 10.1186/s40462-019-0148-6.

[70] Gerhard J.D.M. Müskens, Péter Szinai, Tamas Sapi, Andrea Kölzsch, Martin Wikelski, and Bart A. Nolet. Data from: Flyway connectivity and exchange primarily driven by moult migration in geese [Pannonic population], 2019. URL 10.5441/001/1.46b0mq21.

[71] Rob Slotow, Maria Thaker, and Abi Tamim Vanak. Data from: Fine-scale tracking of ambient temperature and movement reveals shuttling behavior of elephants to water, 2019. URL 10.5441/001/1.403h24q5.

[72] Maria Thaker, Pratik R. Gupte, Herbert H. T. Prins, Rob Slotow, and Abi T. Vanak. Fine-Scale Tracking of Ambient Temperature and Movement Reveals Shuttling Behavior of Elephants to Water. Frontiers in Ecology and Evolution, 7, jan 2019. doi: 10.3389/fevo.2019.00004. URL 10.3389/fevo.2019.00004.

[73] Mark Hebblewhite, Evelyn H. Merrill, Hans Martin, Jodi E. Berg, Holger Bohm, and Scott L. Eggeman. Data from: Study “Ya Ha Tinda elk project, Banff National Park, 2001-2020 (females)”, 2020. URL 10.5441/001/1.5g4h5t6c.

[74] Marlee A. Tucker, Katrin Böhning-Gaese, William F. Fagan, John M. Fryxell, Bram Van Moorter, Susan C. Alberts, Abdullahi H. Ali, Andrew M. Allen, Nina Attias, Tal Avgar, Hattie Bartlam-Brooks, Buuveibaatar Bayarbaatar, Jerrold L. Belant, Alessandra Bertassoni, Dean Beyer, Laura Bidner, Floris M. van Beest, Stephen Blake, Niels Blaum, Chloe Bracis, Danielle Brown, P. J. Nico de Bruyn, Francesca Cagnacci, Justin M. Calabrese, Constança Camilo-Alves, Simon Chamaillé-Jammes, Andre Chiaradia, Sarah C. Davidson, Todd Dennis, Stephen DeStefano, Duane Diefenbach, Iain Douglas-Hamilton, Julian Fennessy, Claudia Fichtel, Wolfgang Fiedler, Christina Fischer, Ilya Fischhoff, Christen H. Fleming, Adam T. Ford, Susanne A. Fritz, Benedikt Gehr, Jacob R. Goheen, Eliezer Gurarie, Mark Hebblewhite, Marco Heurich, A. J. Mark Hewison, Christian Hof, Edward Hurme, Lynne A. Isbell, René Janssen, Florian Jeltsch, Petra Kaczensky, Adam Kane, Peter M. Kappeler, Matthew Kauffman, Roland Kays, Duncan Kimuyu, Flavia Koch, Bart Kranstauber, Scott LaPoint, Peter Leimgruber, John D. C. Linnell, Pascual López-López, A. Catherine Markham, Jenny Mattisson, Emilia Patricia Medici, Ugo Mellone, Evelyn Merrill, Guilherme de Miranda Mourão, Ronaldo G. Morato, Nicolas Morellet, Thomas A. Morrison, Samuel L. Díaz-Muñoz, Atle Mysterud, Dejid Nandintsetseg, Ran Nathan, Aidin Niamir, John Odden, Robert B. O’Hara, Luiz Gustavo R. Oliveira-Santos, Kirk A. Olson, Bruce D. Patterson, Rogerio Cunha de Paula, Luca Pedrotti, Björn Reineking, Martin Rimmler, Tracey L. Rogers, Christer Moe Rolandsen, Christopher S. Rosenberry, Daniel I. Rubenstein, Kamran Safi, Sonia Saïd, Nir Sapir, Hall Sawyer, Niels Martin Schmidt, Nuria Selva, Agnieszka Sergiel, Enkhtuvshin Shiilegdamba, João Paulo Silva, Navinder Singh, Erling J. Solberg, Orr Spiegel, Olav Strand, Siva Sundaresan, Wiebke Ullmann, Ulrich Voigt, Jake Wall, David Wattles, Martin Wikelski, Christopher C. Wilmers, John W. Wilson, George Wittemyer, Filip Zięba, Tomasz Zwijacz-Kozica, and Thomas Mueller. Moving in the Anthropocene: Global reductions in terrestrial mammalian movements. Science, 6374(6374):466–469, jan 2018. doi: 10.1126/science.aam9712. URL 10.1126/science.aam9712.

[75] Scott L. Eggeman, Mark Hebblewhite, Holger Bohm, Jesse Whittington, and Evelyn H. Merrill. Behavioural flexibility in migratory behaviour in a long-lived large herbivore. Journal of Animal Ecology, 3(3):785–797, mar 2016. doi: 10.1111/1365-2656.12495. URL 10.1111/1365-2656.12495.

[76] MARK Hebblewhite, EVELYN H. Merrill, LUIGI E. MORGAN- Tini, CLIFFORD A. White, JAMES R. Allen, ELDON Bruns, LINDA Thurston, and TOMAS E. Hurd. Is the migratory behavior of montane elk herds in peril? the case of alberta’s ya ha tinda elk herd. Wildlife Society Bulletin, 5(5):1280–1294, ec 2006. doi: 10.2193/0091-7648(2006)34[1280:itmbom]2.0.co;2. URL 10.2193/0091-7648(2006)34[1280:ITMBOM]2.0.CO;2.

[77] Mark Hebblewhite and Evelyn H. Merrill. Trade-offs between predation risk and forage differ between migrant strategies in a migratory ungulate. Ecology, 12(12):3445–3454, ec 2009. doi: 10.1890/08-2090.1. URL 10.1890/08-2090.1.

[78] Mark Hebblewhite, Evelyn Merrill, and Greg McDermid. A MULTISCALE TEST OF THE FORAGE MATURATION HYPOTHESIS IN a PARTIALLY MIGRATORY UNGULATE POPULATION. Ecological Monographs, 2(2):141–166, may 2008. doi: 10.1890/06-1708.1. URL 10.1890/06-1708.1.

[79] Ben Koks, Almut Schlaich, Tonio Schaub, Raymond Klaassen, Anny Anselin, Peter Desmet, Tanja Milotic, Kjell Janssens, and Willem Bouten. H_GRONINGEN - Western marsh harriers (Circus aeruginosus, Accipitridae) breeding in Groningen (the Netherlands), 2022. URL 10.5281/zenodo.3552507.

[80] Mark S. Boyce and Simone Ciuti. Data from: Human selection of elk behavioural traits in a landscape of fear, 2020. URL 10.5441/001/1.j484vk24.

[81] Dale G. Paton, Simone Ciuti, Michael Quinn, and Mark S. Boyce. Hunting exacerbates the response to human disturbance in large herbivores while migrating through a road network. Ecosphere, 8(6), jun 2017. doi: 10.1002/ecs2.1841. URL 10.1002/ecs2.1841.

[82] Christina M. Prokopenko, Mark S. Boyce, and Tal Avgar. Characterizing wildlife behavioural responses to roads using integrated step selection analysis. Journal of Applied Ecology, 2(2):470–479, sep 2016. doi: 10.1111/1365-2664.12768. URL 10.1111/1365-2664.12768.

[83] Christina M. Prokopenko, Mark S. Boyce, and Tal Avgar. Extent-dependent habitat selection in a migratory large herbivore: road avoidance across scales. Landscape Ecology, 2(2):313–325, oct 2016. doi: 10.1007/s10980-016-0451-1. URL 10.1007/s10980-016-0451-1.

[84] David R. Roberts, Volker Bahn, Simone Ciuti, Mark S. Boyce, Jane Elith, Gurutzeta Guillera-Arroita, Severin Hauenstein, José J. Lahoz-Monfort, Boris Schröder, Wilfried Thuiller, David I. Warton, Brendan A. Wintle, Florian Hartig, and Carsten F. Dormann. Cross-validation strategies for data with temporal, spatial, hierarchical, or phylogenetic structure. Ecography, 8(8):913–929, mar 2017. doi: 10.1111/ecog.02881 10.1111/ecog.02881.

[85] Henrik Thurfjell, Simone Ciuti, and Mark S. Boyce. Learning from the mistakes of others: How female elk (Cervus elaphus) adjust behaviour with age to avoid hunters. PLOS ONE, 6(6):e0178082, jun 2017. doi: 10.1371/journal.pone.0178082. URL 10.1371/journal.pone.0178082.

[86] Robin A. Benz, Mark S. Boyce, Henrik Thurfjell, Dale G. Paton, Marco Musiani, Carsten F. Dormann, and Simone Ciuti. Dispersal Ecology Informs Design of Large-Scale Wildlife Corridors. PLOS ONE, 11(9): e0162989, sep 2016. doi: 10.1371/journal.pone.0162989. URL 10.1371/journal.pone.0162989.

[87] Erik P. Ensing, Simone Ciuti, Freek A. L. M. de Wijs, Dennis H. Lentferink, Andréten Hoedt, Mark S. Boyce, and Roelof A. Hut. GPS based daily activity patterns in european red deer and north american elk (cervus elaphus): Indication for a weak circadian clock in ungulates. PLoS ONE, 9(9):e106997, sep 2014. doi: 10.1371/journal.pone.0106997. URL 10.1371/journal.pone.0106997.

[88] Joshua Killeen, Henrik Thurfjell, Simone Ciuti, Dale Paton, Marco Musiani, and Mark S Boyce. Habitat selection during ungulate dispersal and exploratory movement at broad and fine scale with implications for conservation management. Movement Ecology, 2(1), jul 2014. doi: 10.1186/s40462-014-0015-4. URL 10.1186/s40462-014-0015-4.

[89] Henrik Thurfjell, Simone Ciuti, and Mark S Boyce. Applications of step-selection functions in ecology and conservation. Movement Ecology, 2(1), feb 2014. doi: 10.1186/2051-3933-2-4. URL 10.1186/2051-3933-2-4.

[90] Simone Ciuti, Tyler B. Muhly, Dale G. Paton, Allan D. McDevitt, Marco Musiani, and Mark S. Boyce. Human selection of elk behavioural traits in a landscape of fear. Proceedings of the Royal Society B: Biological Sciences, 1746(1746):4407–4416, sep 2012. doi: 10.1098/rspb.2012.1483. URL 10.1098/rspb.2012.1483.

[91] Simone Ciuti, Joseph M. Northrup, Tyler B. Muhly, Silvia Simi, Marco Musiani, Justin A. Pitt, and Mark S. Boyce. Effects of Humans on Behaviour of Wildlife Exceed Those of Natural Predators in a Landscape of Fear. PLoS ONE, 11(11):e50611, nov 2012. doi: 10.1371/journal.pone.0050611. URL 10.1371/journal.pone.0050611.

[92] Geert Spanoghe, Peter Desmet, Tanja Milotic, Kjell Janssens, Nico De Regge, Joost Vanoverbeke, and Willem Bouten. MH_ANTWERPEN - Western marsh harriers (Circus aeruginosus, Accipitridae) breeding near Antwerp (Belgium), 2022. URL 10.5281/zenodo.3550093.

[93] Eric W.M. Stienen, Peter Desmet, Tanja Milotic, Francisco Hernandez, Klaas Deneudt, Willem Bouten, Wendt Müller, Hans Matheve, and Luc Lens. LBBG_ZEEBRUGGE - Lesser black-backed gulls (Larus fuscus, Laridae) breeding at the southern North Sea coast (Belgium and the Netherlands), 2022. URL 10.5281/zenodo.3540799.

[94] Eric W.M. Stienen, Peter Desmet, Tanja Milotic, Francisco Hernandez, Klaas Deneudt, Hans Matheve, and Willem Bouten. HG_OOSTENDE - Herring gulls (Larus argentatus, Laridae) breeding at the southern North Sea coast (Belgium), 2022. URL 10.5281/zenodo.3541811.

[95] Rascha J.M. Nuijten, Theo Gerrits, Peter P. De Vries, Gerhard J.D.M. Müskens, and Bart A. Nolet. Data from: Less is more: on-board lossy compression of accelerometer data increases biologging capacity, 2020. URL 10.5441/001/1.8ms7mm80.

[96] Rascha J. M. Nuijten, Theo Gerrits, Judy Shamoun-Baranes, and Bart A. Nolet. Less is more: On-board lossy compression of accelerometer data increases biologging capacity. Journal of Animal Ecology, 1(1):237–247, jan 2020. doi: 10.1111/1365-2656.13164. URL 10.1111/1365-2656.13164.

[97] Andrea Kölzsch, Gerhard J.D.M. Müskens, Peter Glazov, Helmut Kruckenberg, and Martin Wikelski. Data from: Goose parents lead migration V, 2020. URL 10.5441/001/1.ms87s2m6.

[98] A. Kölzsch, A. Flack, G. J. D. M. Müskens, H. Kruckenberg, P. Glazov, and M. Wikelski. Goose parents lead migration V. Journal of Avian Biology, 51(3), mar 2020. doi: 10.1111/jav.02392. URL 10.1111/jav.02392.

[99] Robert A. Ronconi and Katherine R. Shlepr. Data from: Study “Herring Gulls (Larus Argentatus); Ronconi; Brier Island, Canada”, 2020. URL 10.5441/001/1.282vr7kd.

[100] Christine M. Anderson, H. Grant Gilchrist, Robert A. Ronconi, Katherine R. Shlepr, Daniel E. Clark, David A. Fifield, Gregory J. Robertson, and Mark L. Mallory. Both short and long distance migrants use energy-minimizing migration strategies in North American herring gulls. Movement Ecology, 8(1), jun 2020. doi: 10.1186/s40462-020-00207-9. URL 10.1186/s40462-020-00207-9.

[101] Daniel E. Clark, Stuart A. Mackenzie, Kiana Koenen, Jillian Whitney, and Stephen DeStefano. Data from: Study “Herring Gulls (Larus Argentatus); Clark; Massachussets, United States”, 2020. URL 10.5441/001/1.3th8b5q3.

[102] Robert A. Ronconi and Philip D. Taylor. Data from: Study “Herring Gulls (Larus Argentatus); Ronconi; Sable Island, Canada”, 2020. URL 10.5441/001/1.3264ss3v.

[103] Katherine Mertes, Walter Jetz, and Martin Wikelski. Data from: Hierarchical multi-grain models improve descriptions of species’ environmental associations, distribution, and abundance, 2020. URL 10.5441/001/1.cp97k9j1.

[104] Katherine Mertes, Marta A. Jarzyna, and Walter Jetz. Hierarchical multigrain models improve descriptions of species’ environmental associations, distribution, and abundance. Ecological Applications, 30(6), may 2020. doi: 10.1002/eap.2117. URL 10.1002/eap.2117.

[105] Andy M Ramey, S.A. Hatch, Christina A Ahlstrom, M.L. Van Toor, H. Woksepp, J.C. Chandler, John A Reed, Andrew Reeves, J. Waldenstrom, A.B. Franklin, J. Bonnedahl, V.A. Gill, D.M. Mulcahy, and David C Douglas. Tracking Data for Three Large-bodied Gull Species and Hybrids (Larus spp.), 2020. URL 10.5066/P9FZ4OJW.

[106] Geert Spanoghe, Peter Desmet, Tanja Milotic, Gunther Van Ryckegem, Joost Vanoverbeke, Bruno J. Ens, and Willem Bouten. O_WESTERSCHELDE - Eurasian oystercatchers (Haematopus ostralegus, Haematopodidae) breeding in East Flanders (Belgium), 2022. URL 10.5281/zenodo.3734898.

[107] Eric W.M. Stienen, Wendt Müller, Luc Lens, Tanja Milotic, and Peter Desmet. LBBG_JUVENILE - Juvenile lesser black-backed gulls (Larus fuscus, Laridae) hatched in Zeebrugge (Belgium), 2022. URL 10.5281/zenodo.5075868.

[108] R. John Power and Stephen Dell. Data from: A note on the reestablishment of the cheetah population in the Pilanesberg National Park, South Africa, 2020. URL 10.5441/001/1.k6b630mv.

[109] R. John Power, Vincent Van der Merwe, Samantha Page-Nicholson, Mia V. Botha, Stephen Dell, and Pieter Nel. A Note on the Reestablishment of the Cheetah Population in the Pilanesberg National Park, South Africa. African Journal of Wildlife Research, 49(1), feb 2019. doi: 10.3957/056.049.0012. URL 10.3957/056.049.0012.

[110] Geert Spanoghe, Kjell Janssens, Raymond Klaassen, Tonio Schaub, Tanja Milotic, and Peter Desmet. BOP_RODENT - Rodent specialized birds of prey (Circus, Asio, Buteo) in Flanders (Belgium), 2022. URL 10.5281/zenodo.5735405.

[111] Philipp Schwemmer and Stefan Garthe. Data from: Migrating curlews on schedule: departure and arrival patterns of a long-distance migrant depend on time and breeding location rather than on wind conditions, 2021. URL 10.5441/001/1.715k46g2.

[112] Philipp Schwemmer, Moritz Mercker, Klaus Heinrich Vanselow, Pierrick Bocher, and Stefan Garthe. Migrating curlews on schedule: departure and arrival patterns of a long-distance migrant depend on time and breeding location rather than on wind conditions. Movement Ecology, 9(1), mar 2021. doi: 10.1186/s40462-021-00252-y. URL 10.1186/s40462-021-00252-y.

[113] Ben Carlson, Shay Rotics, Ran Nathan, Martin Wikelski, and Walter Jetz. Data from: Individual environmental niches in mobile organisms, 2021. URL 10.5441/001/1.rj21g1p1.

[114] Ben S. Carlson, Shay Rotics, Ran Nathan, Martin Wikelski, and Walter Jetz. Individual environmental niches in mobile organisms. Nature Communications, 12(1), jul 2021. doi: 10.1038/s41467-021-24826-x. URL 10.1038/s41467-021-24826-x.

[115] Antti Piironen, Antti Paasivaara, and Toni Laaksonen. Data from: Birds of three worlds: moult migration to high Arctic expands a boreal-temperate flyway to a third biome, 2021. URL 10.5441/001/1.22kk5126.

[116] Antti Piironen, Antti Paasivaara, and Toni Laaksonen. Birds of three worlds: moult migration to high Arctic expands a boreal-temperate flyway to a third biome. Movement Ecology, 9(1), sep 2021. doi: 10.1186/s40462-021-00284-4. URL 10.1186/s40462-021-00284-4.

[117] Mathieu Basille, Rena R. Borkhataria, A. Lawrence Bryan, David N. Bucklin, Simona Picardi, and Peter C. Frederick. Data from: Study “Wood stork (Mycteria americana) Southeastern US 2004–2019”, 2021. URL 10.5441/001/1.r0h6725k.

[118] Simona Picardi, Peter C. Frederick, Rena R. Borkhataria, and Mathieu Basille. Partial migration in a subtropical wading bird in the southeastern United States. Ecosphere, 11(2), feb 2020. doi: 10.1002/ecs2.3054. URL 10.1002/ecs2.3054.

[119] Adriaan M. Dokter, Kees Oosterbeek, Martin J. Baptist, Peter Desmet, Henk-Jan van der Kolk, Willem Bouten, and Bruno J. Ens. O_BALGZAND - Eurasian oystercatchers (Haematopus ostralegus, Haematopodidae) wintering on Balgzand (the Netherlands), 2022. URL 10.5281/zenodo.5653441.

[120] Kees Oosterbeek, Roeland A. Bom, Judy Shamoun-Baranes, Peter Desmet, Henk-Jan van der Kolk, Willem Bouten, and Bruno J. Ens. O_SCHIERMONNIKOOG - Eurasian oystercatchers (Haematopus ostralegus, Haematopodidae) breeding on Schiermonnikoog (the Netherlands), 2022. URL 10.5281/zenodo.5653477.

[121] Henk-Jan van der Kolk, Kees Oosterbeek, Eelke Jongejans, Magali Frauendorf, Andrew M. Allen, Willem Bouten, Peter Desmet, Hans de Kroon, Bruno J. Ens, and Martijn van de Pol. O_VLIELAND - Eurasian oystercatchers (Haematopus ostralegus, Haematopodidae) breeding and wintering on Vlieland (the Netherlands), 2022. URL 10.5281/zenodo.5653890.

[122] Kees Oosterbeek, Jan de Jong, Peter Desmet, Henk-Jan van der Kolk, Willem Bouten, and Bruno J. Ens. O_AMELAND - Eurasian oystercatchers (Haematopus ostralegus, Haematopodidae) breeding on Ameland (the Netherlands), 2022. URL 10.5281/zenodo.5647596.

[123] Graeme C. Hays, Jeanne A. Mortimer, Alex Rattray, Takahiro Shimada, and Nicole Esteban. Data from: High accuracy tracking reveals how small conservation areas can protect marine megafauna, 2021. URL 10.5441/001/1.r72ph75f.

[124] Graeme C. Hays, Jeanne A. Mortimer, Alex Rattray, Takahiro Shimada, and Nicole Esteban. Data from: High accuracy tracking reveals how small conservation areas can protect marine megafauna. Ecological Applications, 31(7), aug 2021. doi: 10.1002/eap.2418. URL 10.1002/eap.2418.

[125] Mariëlle L. Van Toor, Sergey Kharitonov, Saulius Švažas, Mindaugas Dagys, Eric Kleyheeg, Gerard Müskens, Ulf Ottosson, Ramunas Žydelis, and Jonas Waldenström. Data from: Study “Eurasian wigeon (Mareca penelope) Netherlands Lithuania 2018-2019”, 2021. URL 10.5441/001/1.dv5mm289.

[126] Mariëlle L. van Toor, Sergey Kharitonov, Saulius Švažas, Mindaugas Dagys, Erik Kleyheeg, Gerard Müskens, Ulf Ottosson, Ramunas Žydelis, and Jonas Waldenström. Migration distance affects how closely Eurasian wigeons follow spring phenology during migration. Movement Ecology, 9(1), ec 2021. doi: 10.1186/s40462-021-00296-0. URL 10.1186/s40462-021-00296-0.

[127] Martin Wikelski. MPIAB PNIC hurricane frigate tracking, 2016. Movebank study 6770990 (accessed on 15 February 2022).

[128] Martin Wikelski, Ran Nathan, and Shay Rotics. HUJ MPIAB White Stork GSM E-Obs, 2015. Movebank study 7002955 (accessed on 15 February 2022).

[129] Stephen Blake, Randy Arndt, and Doug Ladd. Dunn Ranch Bison Tracking Project, 2017. Movebank study 8019591 (accessed on 15 February 2022).

[130] Roland Kays. LifeTrack - Great Egrets, 2017. Movebank study 8849813 (accessed on 15 February 2022).

[131] Martin Wikelski, Ran Nathan, and Shay Rotics. HUJ MPIAB White Stork E-Obs, 2022. Movebank study 8863543 (accessed on 15 February 2022).

[132] Martin Wikelski. LifeTrack White Stork Uzbekistan, 2021. Movebank study 9493881 (accessed on 15 February 2022).

[133] H. Azafzaf, C. Feltrup-Azafzaf, A. Flack, M. Wikelski, and W. Fiedler. LifeTrack White Stork Tunisia, 2015. Movebank study 10157679 (accessed on 15 February 2022).

[134] Barb Jensen. Pandion haliaetus Osprey - SouthEast Michigan, 2018. Movebank study 10204361 (accessed on 15 February 2022).

[135] Martin Wikelski. LifeTrack White Stork Loburg, 2022. Movebank study 10449318 (accessed on 15 February 2022).

[136] Martin Wikelski, Ran Nathan, and Shay Rotics. HUJ MPIAB White Stork GSM 2013, 2022. Movebank study 10449698 (accessed on 15 February 2022).

[137] David Barber. Hooded Vulture Africa, 2022. Movebank study 14671003 (accessed on 15 February 2022).

[138] Stefan Garthe. FTZ Geese Wadden Sea, 2022. Movebank study 69724677 (accessed on 15 February 2022).

[139] Dmitrijs Boiko. LifeTrack Whooper Swan Latvia, 2017. Movebank study 92261778 (accessed on 15 February 2022).

[140] Wolfgang Fiedler. LifeTrack Ducks Lake Constance, 2021. Movebank study 236953686 (accessed on 15 February 2022).

[141] Ryan Askren. Canada geese (Branta canadensis), 2022. Movebank study 329155299 (accessed on 15 February 2022).

[142] PaweŁ Mirski. White-tailed Eagle Poland., 2015. Movebank study 384868221 (accessed on 15 February 2022).

[143] Laurel Serieys. Coyote Valley Bobcat Habitat Connectivity Study, 2018. Movebank study 475878514 (accessed on 15 February 2022).

[144] Laurel Serieys. Aromas Hills Bobcat Habitat Connectivity Study, 2018. Movebank study 501787846 (accessed on 15 February 2022).

[145] Harald Grabenhofer. Graugans Zugverhalten Neusiedler See, 2022. Movebank study 505156776 (accessed on 15 February 2022).

[146] Patrick Scherler. Milvus_milvus_atlantismarcuard, 2021. Movebank study 672882373 (accessed on 15 February 2022).

[147] James P. Lawler. NPS Dall Sheep in Yukon-Charley Rivers National Preserve, 2003. Movebank study 673728219 (accessed on 15 February 2022).

[148] Perilhon. Milvus migrans, 2022. Movebank study 892924356 (accessed on 15 February 2022).

[149] Thierry Boulinier. Ecopath, Brown skua, Boulinier et al., Amsterdam Island, 2020. Movebank study 918219824 (accessed on 15 February 2022).

[150] Frédéric Jiguet. Birdman research project. Tracking curlews to unravel migration connectivity., 2022. Movebank study 1077731101 (accessed on 15 February 2022).

[151] Scott LaPoint. Carnivore movements near Black Rock Forest New York, 2021. Movebank study 1088836380 (accessed on 15 February 2022).

[152] Chris H. Fleming. GPS calibration data (global), 2019. Movebank study 1092737859 (accessed on 15 February 2022).

[153] Morten Frederiksen. Ivory gull N Greenland 2018/19, 2023. Movebank study 1123149708 (accessed on 15 February 2022).

[154] INTERREX. Caspian Gulls - Poland, 2020. Movebank study 1208105916 (accessed on 15 February 2022).

[155] Petras Kurlavičius. Common Crane 2020 (Lithuanian University of Educational Studies; LEU), 2022. Movebank study 1229945587 (accessed on 15 February 2022).

[156] Dominique Berteaux. Arctic fox Bylot - GPS tracking, 2021. Movebank study 1241071371 (accessed on 15 February 2022).

[157] Frédéric Jiguet. Corvus corone [ID_PROG 883], 2022. Movebank study 1266784970 (accessed on 15 February 2022).

[158] Jérôme Cavailhes. Monitoring of Capra ibex (Bovidae) populations in the western alps (project ALCOTRA LEMED-IBEX), 2020. Movebank study 1285079529 (accessed on 15 February 2022).

[159] Martin Wikelski. Cathartes aura MPIAB Cuba, 2022. Movebank study 1393954358 (accessed on 15 February 2022).

[160] Jelle Loonstra. Lapwing NFW Vanellus Vanellus, 2022. Movebank study 1448409403 (accessed on 15 February 2022).

[161] Sinan Robillard. Variability of White Stork flight patterns prior to earth-quakes, 2021. Movebank study 1498452485 (accessed on 15 February 2022).

[162] Wolfgang Fiedler. LifeTrack White Stork Sarralbe [ID_PROG 1093], 2022. Movebank study 1562253659 (accessed on 15 February 2022).

[163] Willem Burger. Tchad Redneck Ostrich, 2022. Movebank study 1671751878 (accessed on 15 February 2022).

[164] Patricia Medici. Lowland tapirs, Tapirus terrestris, in Southern Brazil, 2019. Movebank study 1907973121 (accessed on 15 February 2022).

[165] Diana Solovyeva. Vega gull (Larus vegae) - GPS - Russia South Korea Japan, 2019. Movebank study 1907974323 (accessed on 15 February 2022).

[166] Simon Chamaillé-Jammes. Plains zebra Chamaillé-Jammes Hwange NP, 2016. Movebank study 295134472.

[167] Arnold Tshipa, Hugo Valls-Fox, Hervé Fritz, Kai Collins, Lovelater Sebele, Peter Mundy, and Simon Chamaillé-Jammes. Partial migration links local surface-water management to large-scale elephant conservation in the world’s largest transfrontier conservation area. Biological Conservation, 215:46–50, 2017.

[168] Ikuya Yamada, Akari Asai, Jin Sakuma, Hiroyuki Shindo, Hideaki Takeda, Yoshiyasu Takefuji, and Yuji Matsumoto. Wikipedia2Vec: An Efficient Toolkit for Learning and Visualizing the Embeddings of Words and Entities from Wikipedia. In Proceedings of the 2020 Conference on Empirical Methods in Natural Language Processing: System Demonstrations, pages 23–30, Online, October 2020. Association for Computational Linguistics. doi: 10.18653/v1/2020.emnlp-demos.4. URL https://aclanthology.org/2020.emnlp-demos.4.

[169] Oscar Venter, Eric W. Sanderson, Ainhoa Magrach, James R. Allan, Jutta Beher, Kendall R. Jones, Hugh P. Possingham, William F. Laurance, Peter Wood, Balázs M. Fekete, Marc A. Levy, and James E. M. Watson. Sixteen years of change in the global terrestrial human footprint and implications for biodiversity conservation. Nature Communications, 7(1): 12558, August 2016. ISSN 2041-1723. doi: 10.1038/ncomms12558. URL https://www.nature.com/articles/ncomms12558. Number: 1 Publisher: Nature Publishing Group.

[170] O. Venter, E.W. Sanderson, A. Magrach, J.R. Allan, J. Beher, K.R. Jones, M.A. Levy, and J.E. Watson. Last of the Wild Project, Version 3 (LWP-3): 2009 Human Footprint, 2018 Release, 2018. URL https://sedac.ciesin.columbia.edu/data/set/wildareas-v3-2009-human-footprint. xType: dataset.

[171] Stephen E. Fick and Robert J. Hijmans. WorldClim 2: new 1-km spatial resolution climate surfaces for global land areas. International Journal of Climatology, 12(12):4302–4315, 2017. ISSN 1097-0088. doi: 10.1002/joc.5086. URL https://onlinelibrary.wiley.com/doi/abs/10.1002/joc.5086. _eprint: https://onlinelibrary.wiley.com/doi/pdf/10.1002/joc.5086.

[172] Marcel Buchhorn, Bruno Smets, Luc Bertels, Bert De Roo, Myroslava Lesiv, Nandin-Erdene Tsendbazar, Martin Herold, and Steffen Fritz. Copernicus Global Land Service: Land Cover 100m: collection 3: epoch 2015: Globe, September 2020. URL https://zenodo.org/record/3939038. Type: dataset.

[173] Ashish Vaswani, Noam Shazeer, Niki Parmar, Jakob Uszkoreit, Llion Jones, Aidan N Gomez, Łukasz Kaiser, and Illia Polosukhin. Attention is All you Need. In Advances in Neural Information Processing Systems, volume 30. Curran Associates, Inc., 2017. URL https://proceedings.neurips.cc/paper/2017/hash/3f5ee243547dee91fbd053c1c4a845aa-Abstract.html.

[174] Cédric Villani. Optimal Transport, volume 338 of Grundlehren der mathematischen Wissenschaften. Springer Berlin Heidelberg, Berlin, Heidelberg, 2009. ISBN 9783540710493 9783540710509. doi: 10.1007/978-3-540-71050-9. URL http://link.springer.com/10.1007/978-3-540-71050-9.

[175] Dan Hendrycks and Kevin Gimpel. Gaussian Error Linear Units (GELUs), July 2020. URL http://arxiv.org/abs/1606.08415. xarXiv:1606.08415 [cs].

[176] Ondřej Cífka and Antoine Liutkus. Black-box language model explanation by context length probing, December 2022. URL http://arxiv.org/abs/2212.14815. xarXiv:2212.14815 [cs].

[177] Leo Breiman. Random Forests. Machine Learning, 1(1):5–32, October 2001. ISSN 1573-0565. doi: 10.1023/A:1010933404324. URL 10.1023/A:1010933404324.

[178] Aaron Fisher, Cynthia Rudin, and Francesca Dominici. All Models are Wrong, but Many are Useful: Learning a Variable’s Importance by Studying an Entire Class of Prediction Models Simultaneously. Journal of Machine Learning Research, 177(177):1–81, 2019. ISSN 1533-7928. URL http://jmlr.org/papers/v20/18-760.html.

[179] Giles Hooker, Lucas Mentch, and Siyu Zhou. Unrestricted permutation forces extrapolation: variable importance requires at least one more model, or there is no free variable importance. Statistics and Computing, 6(6):82, October 2021. ISSN 1573-1375. doi: 10.1007/s11222-021-10057-z. URL 10.1007/s11222-021-10057-z.

[180] Pascal Marchand, Mathieu Garel, Gilles Bourgoin, Antoine Duparc, Dominique Dubray, Daniel Maillard, and Anne Loison. Combining familiarity and landscape features helps break down the barriers between movements and home ranges in a non-territorial large herbivore. Journal of Animal Ecology, 2(2):371–383, 2017. ISSN 1365-2656. doi: 10.1111/1365-2656.12616. URL https://onlinelibrary.wiley.com/doi/abs/10.1111/1365-2656.12616. _eprint: https://besjournals.onlinelibrary.wiley.com/doi/pdf/10.1111/1365-2656.12616.

[181] Marlee A. Tucker, Katrin Böhning-Gaese, William F. Fagan, John M. Fryxell, Bram Van Moorter, Susan C. Alberts, Abdullahi H. Ali, Andrew M. Allen, Nina Attias, and Tal Avgar. Moving in the Anthropocene: Global reductions in terrestrial mammalian movements. Science, 6374(6374):466–469, 2018. Publisher: American Association for the Advancement of Science.

[182] Ondřej Cífka. cifkao/moveformer: 20230805, August 2023. URL 10.5281/zenodo.8217149.

[183] Ondřej Cífka. cifkao/gps2var: v0.1.0-alpha.1, June 2022. URL 10.5281/zenodo.8217156.

[184] Ondřej Cífka, Simon Chamaillé-Jammes, and Antoine Liutkus. Trained models and metrics from: MoveFormer: a Transformer-based model for step-selection animal movement modelling, March 2023. URL 10.5281/zenodo.7698263.

